# Genetic load, eco-evolutionary feedback and extinction in metapopulations

**DOI:** 10.1101/2023.12.02.569702

**Authors:** Oluwafunmilola Olusanya, Ksenia Khudiakova, Himani Sachdeva

## Abstract

Habitat fragmentation poses a significant risk to population survival, causing both demographic stochasticity and genetic drift within local populations to increase, thereby increasing genetic load. Higher load causes population numbers to decline, which reduces the efficiency of selection and further increases load, resulting in a positive feedback which may drive entire populations to extinction. Here, we investigate this eco-evolutionary feedback in a metapopulation consisting of local demes connected via migration, with individuals subject to deleterious mutation at a large number of loci. We first analyse the determinants of load under soft selection, where population sizes are fixed, and then build upon this to understand hard selection, where population sizes and load co-evolve. We show that under soft selection, very little gene flow (less than one migrant per generation) is enough to prevent fixation of deleterious alleles. By contrast, much higher levels of migration are required to mitigate load and prevent extinction when selection is hard, with critical migration thresholds for metapopulation persistence increasing sharply as the genome-wide deleterious mutation rate becomes comparable to the baseline population growth rate. Moreover, critical migration thresholds are highest if deleterious mutations have intermediate selection coefficients, but lower if alleles are predominantly recessive rather than additive (due to more efficient purging of recessive load within local populations). Our analysis is based on a combination of analytical approximations and simulations, allowing for a more comprehensive understanding of the factors influencing load and extinction in fragmented populations.

## Introduction

The long-held belief that ecological and evolutionary processes do not influence each other has been challenged in recent years, with many studies documenting ecological and evolutionary change on contemporaneous timescales (Hairston Jr et al., 2005; Thompson, 1998). This might occur when selection varies sharply over space and time, or when populations are too small to adapt efficiently, resulting in rapid loss of fitness (e.g., due to the accumulation of deleterious alleles), which feeds back into demography.

Such eco-evolutionary feedback is particularly important in fragmented landscapes with many small-sized patches, where both inbreeding (mating between genetically related individuals) and genetic drift (stochastic changes in genetic composition) can reduce the efficiency of natural selection. As a consequence, populations tend to fix more deleterious mutations, causing genetic load to increase and population numbers to decline, which can further exacerbate both genetic drift and inbreeding. The resultant positive feedback between increasing load and declining population sizes may drive populations towards extinction through a process termed *mutational meltdown* (Lynch et al., 1995b). Evidence of such extinction events has been found in plants (Matthies et al., 2004), vertebrates (Fagan and Holmes, 2006) and invertebrates (Saccheri et al., 1998). This kind of feedback is further aggravated by demographic and environmental stochasticity, both of which hinder adaptation to changing environments.

Migration between local populations in a fragmented landscape can affect various aspects of this ecoevo feedback loop, homogenizing both genetic composition and population density across the population. Investigations of natural populations yield examples of both the beneficial and harmful effects of migration. For instance, Finger et al. (2011) demonstrated how augmented gene flow reduced inbreeding in an isolated population of the jellyfish tree. Similarly, Land and Lacy (2000) showed that introducing eight wild-caught Texan females into a small isolated Florida panther population tripled its size, bringing it back from near extinction within just twelve years. On the flip side, there are also several examples of hybridisation-induced extinction, e.g., due to gene flow between domesticated species and their wild relatives (Todesco et al., 2016). These examples thus highlight the diverse effects of migration– ranging from ‘rescue’ (e.g., when migrants are introduced into genetically depauperate populations) to ‘swamping’ (e.g., due to gene flow between populations adapted to different environments), and more generally, the tension between inbreeding and outbreeding depression (Edmands, 2007; Frankham et al., 2011; Templeton, 1986). This introduces complexities to practical decisions in conservation such as those concerning assisted gene flow (Aitken and Whitlock, 2013), underscoring the need for a more quantitative and theoretical understanding of gene flow in fragmented populations.

Migration can have complex effects on genetic diversity even in populations under *spatially uniform* selection, e.g., when all local populations are subject to purifying selection against the same unconditionally deleterious alleles. Different local populations may be nearly fixed for deleterious alleles at different loci just by chance. Migration between them will then result in hybrid offspring of increased vigour in which recessive deleterious mutations are masked. Migration thus increases fitness variation within local demes, which has two competing effects– it prevents fixation of deleterious alleles and alleviates drift load (especially at low levels of migration), but also hinders purging of recessive alleles by increasing heterozygosity (at higher levels of migration) (Glémin et al., 2003).

The effect of migration on genetic diversity and adaptation in structured populations depends crucially on whether selection is ‘soft’ or ‘hard’, i.e., whether different local populations contribute equally to the next generation regardless of fitness, or if the contribution of fitter populations is higher (e.g. when these are larger and send out more migrants). However, most work on hard vs. soft selection in subdivided populations assumes a rather specialised life cycle in which all individuals from all demes join a common mating pool prior to reproduction, followed by uniform redistribution of offspring back into demes (Christiansen, 1975; Levene, 1953; Ravigné et al., 2004), thus not allowing for the more realistic possibility of *limited* dispersal between demes. Moreover, models of hard selection implicitly assume that the metapopulation as a whole is under global density regulation and at carrying capacity (see, e.g., Whitlock (2002)). Thus, such models cannot be used to understand when and how the positive feedback between declining population size and increasing load might lead to local or global extinction, and to what extent this may be arrested by (limited) gene flow between populations. This therefore calls for more realistic models that allow for variation in both local and global population sizes and explicitly incorporate eco-evolutionary feedback.

Lynch et al.(1995a; 1995b) considered such a model, where deleterious mutation accumulation can lead to extinction. Using extensive computer simulations, they showed that populations with low effective size (typically *<*100) are at significant risk of extinction via mutational meltdown within only about 100 generations (Lynch et al., 1995a)– a conclusion with potential implications, e.g., for captive breeding programs. However, their analysis was limited to a single randomly mating population. Higgins and Lynch (2001) generalised this to a metapopulation with global and nearest-neighbour dispersal, and showed that the eventual fate of a metapopulation (as measured by the median time to extinction) depends critically on the number of demes (see also Lande et al. (2003), Ch 4, Table. 4.1) as well as on the dispersal neighbourhood. Their analysis however primarily focused on extinction times and was restricted to scenarios with relatively small patch sizes, resulting in early population extinction. Furthermore, their study was entirely simulation-based, making it hard to generalize their conclusions.

Going beyond simulations, Szép et al. (2021) derived analytical results for local adaptation and extinction in a metapopulation allowing for genetic drift, demographic stochasticity and explicit eco-evo feedback between (local) population size and load. Crucially, for local adaptation involving *polygenic* traits, small shifts in deleterious allele frequency at many loci can substantially increase total load, thus decreasing population size, which further increases drift relative to selection per locus. This thus generates an indirect ‘eco-evolutionary’ coupling between different loci, even in the absence of epistasis or linkage disequilibria. Szép et al. (2021) showed that relatively low levels of gene flow can destroy local adaptation and drive populations to extinction when selection is hard and if adaptive traits are more polygenic.

This begs the question: how does gene flow influence extinction risk in other scenarios, e.g., when selection is uniform across populations, and how are its effects mediated by the mutation target and effect sizes of deleterious alleles? Using a similar theoretical model (but with uniform selection), Sachdeva et al. (2022) investigated whether gene flow from a large mainland can arrest mutational meltdown of an island population. They found that migration has qualitatively different effects depending on the size of the island population– reducing load and the risk of extinction when the population is small and isolated, but increasing load (by hindering purging) when it is large and connected. They also identified the total deleterious mutation rate relative to the population growth rate as a key determinant of extinction.

However, going beyond the special case of marginal populations subject to asymmetric gene flow, it is unclear how eco-evo feedback influences extinction risk and load more generally in fragmented landscapes, where no one local population may be large enough to act as a reservoir of fitness-increasing alleles. Instead, different populations may locally fix for different subsets of deleterious alleles that can potentially mask one another, when connected via migration. In this study, we analyse this scenario in detail by explicitly modeling the eco-evolutionary dynamics of population size and allele frequencies in a metapopulation made up of a large number of islands (demes) exchanging migrants at random. We investigate the effect of gene flow on equilibrium load and population size, as well as the critical migration thresholds required for the persistence (i.e., for non-extinction) of the metapopulation. We analyse how these thresholds are influenced by the architecture of load (i.e., the total deleterious mutation rate and the selection and dominance coefficients of mutations) as well as features of the metapopulation landscape (carrying capacities and baseline growth rate of local demes). As we see below, a key parameter governing the fate of the metapopulation is the total mutation rate relative to the baseline population growth rate, which determines the extent to which genetic load depresses population growth and can thus be considered a proxy for the ‘hardness of selection’. Our analysis shows that much higher levels of migration are required for metapopulation persistence when selection is hard, i.e., the total deleterious mutation rate comparable to the baseline growth rate. However, critical migration thresholds are also shaped by the selection and dominance coefficients of deleterious alleles, being highest for moderately deleterious alleles with additive (co-dominant) effects. In addition to explicitly modeling the direct eco-evo coupling between population size and load, our theoretical approach also accounts for the indirect coupling between the dynamics of different selected loci (see above). This indirect coupling adds complexity to eco-evo dynamics and can have significant implications for population fitness and evolutionary outcomes in the metapopulation.

## Model and Methods

We consider a metapopulation with a large number *D* of demes (local populations) connected via migration. Individuals are diploid and subject to deleterious mutation at *L* unlinked loci at rate *u* per haploid locus, resulting in a genome-wide deleterious mutation rate of 2*Lu*. We also allow for reverse mutations at rate *v*. Fitness is assumed to be multiplicative across loci. At any locus, the wild-type, heterozygote and deleterious homozygote have fitness 1, *e*^−*hs*^ and *e*^−*s*^ respectively, with *h* representing the dominance coefficient and *s*, the strength of selection against the deleterious homozygote. In the main paper, we only consider equal effects (*s* and *h* same) across all loci; this assumption is relaxed in Online Supplement C.

In every generation, a fraction *m* of individuals from each deme migrate into a common pool; individuals from this pool are then uniformly redistributed across all demes. Migration is followed by mutation, which is followed by density and fitness-dependent reproduction. More specifically, any individual *j* in deme *i* produces a Poisson-distributed number of gametes with mean equal to 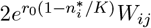, where *W*_*ij*_ is the fitness of the individual (as described above), 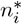 is the population size in deme *i* (after migration), *K* is the carrying capacity and *r*_0_ the baseline growth rate in the deme. Each gamete is created by free recombination between the two parental haplotypes, and gametes paired at random to create diploid offspring, which then replace parents (resulting in discrete, non-overlapping generations). Thus, the population size at the end of reproduction is also Poisson-distributed with mean 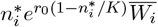 (where 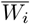 is the mean fitness in deme *i* prior to reproduction), so that population growth is reduced relative to the baseline growth rate *r*_0_ by both genetic load and logistic density-dependence within demes. Note that even though *K, r*_0_, and the deleterious effects of mutations are the same across all demes, population sizes can still differ across demes due to stochastic variation in local population fitness 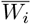 and demographic stochasticity.

Thus, the model parameters are: the number of loci *L*, the per locus mutation rates *u* and *v* (which we assume to be equal), the per locus selection strength *s* and dominance coefficient *h*, the migration rate *m*, the baseline growth rate *r*_0_ and carrying capacity *K* per deme. However, as we demonstrate below, in the continuous time limit, the co-evolution of allele frequencies and population sizes is more conveniently described in terms of scaled (dimensionless) parameters. These parameters are: *Km, Ks* and *Ku* (which represent respectively the number of migrants per deme, the per locus strength of selection relative to drift within a deme, and the average number of deleterious mutations arising per locus per generation per deme, all defined for demes at *carrying capacity*), the dominance coefficient *h*, the genomewide mutation rate scaled by the baseline growth rate (2*Lu*)*/r*_0_ (which is a measure of the ‘hardness’ of selection) and *r*_0_*K* (which determines the strength of demographic fluctuations). The last two parameters warrant further explanation: the expected genetic load in a *large* population at mutation-selection balance is precisely 2*Lu*, regardless of *s* and *h*, if alleles are not completely recessive. Thus, (2*Lu*)*/r*_0_ represents the relative reduction in the growth rate of a large population due to mutation load. Finally, the parameter *r*_0_*K* represents the average number of births (or deaths) in a deme at carrying capacity. Thus, larger values of *r*_0_*K* correspond to weaker demographic stochasticity.

To understand how different evolutionary processes affect population outcomes, we first analyse *soft selection*, which corresponds to the (2*Lu*)*/r*_0_ → 0 limit of the model. Under soft selection, population sizes are constant and the same across demes (*n*_*i*_=*K* for all *i*), regardless of load. We then consider *hard selection*, where genetic load is comparable to the baseline growth rate (i.e., 2*Lu/r*_0_≲1), so that population size decreases with increasing maladaptation. Also, since sizes can vary randomly across demes (due to stochastic variation in allele fitness), larger and more fit demes contribute more to the migrant pool, while also being less influenced by incoming migrants. The hard selection model thus allows for both local extinctions (wherein a subset of demes go into a ‘sink’ state) or global extinction of the metapopulation.

Before analysing the two selection regimes, it is useful to contextualise our use of the term ‘hard selection’, which has been used for two distinct kinds of models with subdivision in previous work. One class of models posits hard selection as a process wherein local population sizes decline with increasing load; these models thus explicitly track the co-evolution of population size and allele frequencies, and allow for extinction (Polechová and Barton, 2015; Szép et al., 2021). By contrast, in the second class of models, hard vs. soft selection refers only to whether or not local populations contribute to the next generation in proportion to their fitness; in these models, population sizes are determined solely by some form of (local vs. global) density-dependent regulation (Christiansen, 1975; Whitlock, 2002). The latter class of models implicitly assume that the population as a whole is at carrying capacity, thus neglecting any influence of genetic load on population size and consequently, on the extent of drift. Our definition of hard selection is consistent with the first class of models, in particular, allowing (sub-)populations to go extinct.

Our analysis utilises simulations as well as analytical approximations. We analyse soft selection using the *diffusion approximation*, which predicts the equilibrium distribution of allele frequencies in any deme conditional on the mean allele frequencies across the full metapopulation (Wright, 1937). In doing this, we assume that alleles at different loci evolve independently-this holds approximately if recombination is much faster than all evolutionary and ecological processes, but becomes less accurate when load is substantial, i.e., *O*(1). We thus study the effects of associations between loci using effective migration rates (Bengtsson, 1985), that can be incorporated into the diffusion approximation (Sachdeva, 2022).

Under hard selection, population sizes can vary between demes and also within a deme over time, requiring us to follow the joint distribution of population sizes and allele frequencies across all demes (Szép et al., 2021). However, it turns out that we cannot explicitly solve for this distribution under the diffusion approximation in the presence of mutation. Hence, to make analytical progress we use the *semi-deterministic approximation* (see also Sachdeva et al., 2022). This assumes that the size of any deme is reduced relative to carrying capacity by an amount that is *determined* by load, rather than following a distribution. It further assumes that the load is that expected at mutation-selection-migration-drift equilibrium for this size, allowing us to solve self-consistently for the population size (and load). As detailed below, this approximation accounts for genetic drift but not demographic stochasticity, and is hence ‘semi-deterministic’.

### Evolution of allele frequencies and population sizes in continuous time

If recombination is much faster than ecological and evolutionary processes, then loci evolve independently, allowing us to track allele (instead of genotype) frequencies. Further, if allele frequencies and population sizes change only slightly per generation, then one can describe their dynamics via *continuous time* equations. Let *p*_*i,j*_ denote the deleterious allele frequency at locus *j* (where *j*=1, 2, … *L*) in deme *i* (where *i*=1, 2, … *D*), *q*_*i,j*_=1−*p*_*i,j*_ the corresponding wildtype frequency, and *n*_*i*_, the population size in deme *i*. Then *{p*_*i,j*_*}* and *n*_*i*_ satisfy the following coupled stochastic differential equations:

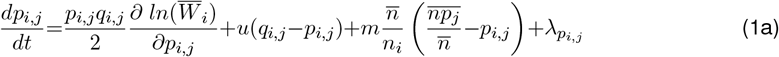

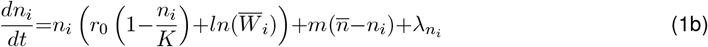

In the above equations, 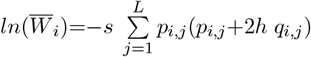 is the mean log fitness in deme *i*; alternatively,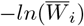 is the genetic load and is obtained by summing over the load contributions across all *L* loci. Further, 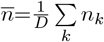 is the mean population size per deme and 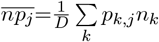 the mean number of deleterious alleles (at locus *j*) per deme, averaged across all demes. Finally, 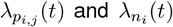 are independent Gaussian white noise processes with mean zero and variances given by 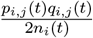 and *n*_*i*_(*t*) respectively. These capture the effects of genetic drift and demographic stochasticity on allele frequencies and population sizes respectively. Note that in our model, the strength of genetic drift and demographic stochasticity both depend on local deme sizes, since density regulation occurs *within* demes.

Equation 1a describes the dynamics of allele frequencies in any deme, with the four terms capturing the effects of selection, mutation, migration and genetic drift on allele frequencies. The third term is noteworthy: it describes how migration pulls allele frequencies within any deme towards 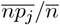, the mean frequency across the metapopulation. However, this effect is modulated by 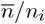, being stronger for islands with less-than-average population sizes which receive more immigrants than they send out emigrants.

Equation 1b describes the dynamics of population size in any deme. The first term captures the effects of fitness-dependent population growth and logistic regulation; in particular, the baseline growth rate of any population is reduced by an amount equal to 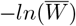, so that demes with higher load have lower growth rates. The second term describes the effects of migration, which tends to pull population sizes towards the metapopulation average, and the third term captures the effects of demographic stochasticity.

### Scaled parameters and separation of timescales

It is convenient to work with scaled parameters, which can be constructed by scaling population size and time by the carrying capacity, i.e., *N* =*n/K* and *T* =*t/K*, and all evolutionary and ecological rates by 1*/K*, so that we have the scaled parameters *Ks, Ku, Km* and *ζ*=*r*_0_*K*. Alternatively, we can scale time as *τ* =*r*_0_*t* and all rates by *r*_0_, the baseline growth rate of populations, thus yielding the scaled parameters *s/r*_0_, *u/r*_0_, *m/r*_0_. The choice of scaling parameters depends on whether one follows dynamics over evolutionary (e.g., local coalescence) or ecological time scales, which are governed respectively by *K* and 1*/r*_0_. Equation (1) can be re-expressed in terms of scaled parameters as:

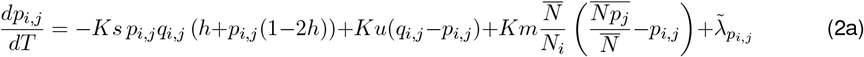

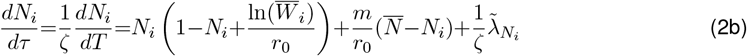

where (as before) 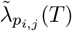 and 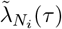 are independent Gaussian white noise processes with mean zero and variances 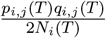 and *N*_*i*_(*τ*) respectively. If *r*_0_≫1*/K* (or *ζ*=*r*_0_*K*≫1), i.e., ecological timescales are much shorter than evolutionary timescales, then the third term in eq. (2b), which captures the effect of demographic stochasticity on population sizes, can be neglected. Further, the second term, which scales as *m/r*_0_ (or equivalently, (*Km*)*/ζ*) is also negligible, provided *Km*, the average number of migrants per deme is *𝒪* (1). Thus, in the limit *ζ*=*r*_0_*K*≫1, population sizes equilibrate much faster than allele frequencies (over a timescale that is essentially governed by 1*/r*_0_) towards a value that is essentially determined by load relative to *r*_0_. This separation of timescales is at the core of our semi-deterministic approximation (details below).

### Diffusion approximation for allele frequencies

Under soft selection, each deme is at carrying capacity, independent of load, so that: 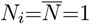 and 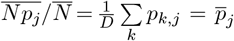 in eq. (2a). Thus, allele frequencies evolve independently of population size (which is constant) and of each other, so that the joint equilibrium distribution Ψ (*p*_1_, *p*_2_, …, *p*_*L*_) is simply the product of the marginal distributions *ψ* (*p*_*j*_). Moreover, if the number of demes is very large (*D*→∞), then following Wright (1937), we can write down an explicit expression for the allele frequency distribution *ψ*(*p*_*j*_) at any locus *j* in any deme, conditioned on the mean allele frequency 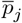 across the full metapopulation (which is also the mean in the migrant pool):

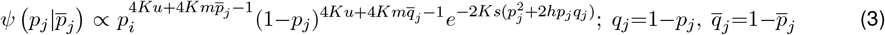

One can numerically integrate over this distribution to obtain the expected frequency 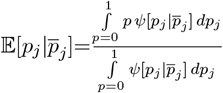 as a function of 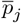. This satisfies 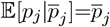, allowing us to numerically solve for 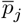, which can then be substituted into eq. (3) to obtain an explicit expression for the allele frequency distribution. Integrating over this distribution now yields the expected heterozygosity, load etc.

### Semi-deterministic approximation for population size and allele frequencies

The diffusion approximation does not yield an explicit solution for the joint distribution of population sizes and allele frequencies (in the presence of mutation). Thus, we employ an approximate *semi-deterministic* analysis which accounts for the effects of genetic drift on allele frequencies, but not those of demographic fluctuations on population size (Mullon and Lehmann, 2018; Sachdeva et al., 2022; Szép et al., 2021). As discussed above, this can be justified when deme sizes are large and *ζ*=*r*_0_*K*≫1, so that the third term in eq. (2b) (which captures the effect of demographic stochasticity) is negligible. Nevertheless, if individual alleles are weakly deleterious, so that *Ks* is *𝒪* (1), then genetic drift still has appreciable effects on allele frequencies. Thus, under the semi-deterministic approximation, we assume that the equilibrium population size in each deme is *determined* by the load, rather than following a distribution. However, allele frequencies are characterised by a distribution that depends on the extent of drift, which in turn depends on local population size.

More formally, we take the limit *ζ*=*r*_0_*K*→∞, so that both the second and third term in eq. (2b) (which scale as 1*/ζ* and (*Km*)*/ζ* respectively) are negligible. Neglecting the second term (which captures the effect of migration on population sizes) is valid as long as *Km*, the number of migrants per deme for demes at carrying capacity, is *𝒪* (1), so that (*Km*)*/ζ* is small. Alternatively, it can be justified in a scenario where every deme has population size close to 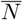, the metapopulation average, so that no deme acts as a demographic source or sink. Then, at equilibrium (i.e., setting *dN/dτ* =0 in eq. (2b)), we have either *N* =0 (local extinction) or 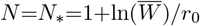 (a ‘stable population’ state). Thus, demes can exist in a stable, non-extinct state (with *N*_*_*>*0) if the genetic load, 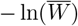, is less than the baseline growth rate *r*_0_, where 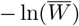 itself depends on *N*_*_. If we further assume that the load is close to the value expected under selection-mutation-migration-drift equilibrium at *N*_*_ (and is thus predicted by eq. (3)), then we have the following coupled equations for population size and mean deleterious allele frequency at equilibrium:

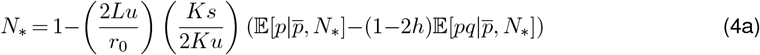

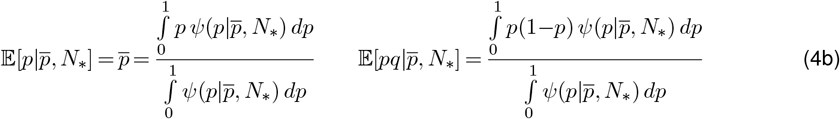

where: 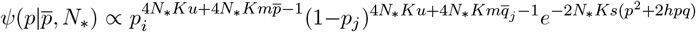

In order to numerically solve equation (4), we start with an initial guess for 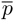 and *N*_*_, use this to compute 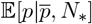 and 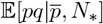 (using eq. (4b)), then use these to compute the new *N*_*_ (using eq. (4a)) as well as the new 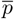 (by setting it equal to 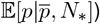), and then iterate until 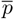 and *N*_*_ converge to fixed values that do not change over successive iterations. The final fixed point can be sensitive to the initial values for 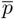 and *N*_*_. Thus, we always start with 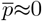 and *N*_*_≈1: this always leads to a fixed point corresponding to a ‘stable population’ equilibrium (with *N*_*_*>*0), provided such an equilibrium exists and is stable.

Note that under the semi-deterministic approximation (as described by equation (4)), population outcomes depend on only five *composite* parameters: the dominance coefficient *h*; the per locus selection strength scaled by the carrying capacity *Ks*, the per locus mutation rate scaled by the carrying capacity *Ku*, the migration rate scaled by the carrying capacity *Km*, and the genomewide mutation rate relative to the baseline growth rate, (2*Lu*)*/r*_0_, that determines the hardness of selection. As we see below in fig. 3, the semi-deterministic approximation becomes increasingly accurate as we take the limit *ζ*=*r*_0_*K*→∞, while holding all scaled parameters *Ks, Ku, Km* and (2*Lu*)*/r*_0_ constant.

Two remarks about the semi-deterministic approximation are in order. First, in parameter regimes where two stable equilibria *N* =0 and *N* =*N*_*_*>*0 are possible, different demes within the metapopulation may reach different equilibria, such that some fraction of demes is nearly extinct, while the remainder support large, stable populations. In such a scenario, we cannot predict the proportion of extinct vs. stable demes, though the semi-deterministic approximation does accurately predict the average population size *N*_*_ in non-extinct demes. Second, where different equilibria are possible, demes may switch from one equilibrium to another, resulting in a turnover between nearly extinct and stable demes in the metapopulation. The key assumption underlying the semi-deterministic approximation is that any individual deme switches very rarely so that population sizes are stable for long periods of time. This allows enough time for allele frequencies to equilibrate at a given *N*_*_, so that genetic load within a deme can be approximated by the expectation at mutation-selection-drift-migration balance.

### Stochastic simulations

Since multi-locus individual-based simulations in metapopulations are computationally expensive, we use these in a targeted way to identify parameter regimes where multi-locus associations have a significant effect on allele frequency distributions under soft selection. It turns out that such associations only have limited effects which are captured reasonably well by incorporating effective migration rates into the diffusion approximation (Figure 2; see also Sachdeva, 2022 and Zwaenepoel et al., 2023). Thus, for the case of hard selection, we only perform allele frequency simulations that neglect associations, i.e., assume linkage and identity equilibrium between loci.

These simulations follow population sizes *n*_*i*_ and allele frequencies *p*_*i,j*_ at all *L* loci in all *D* demes (where *i*=1, …, *D* and *j*=1, …, *L*), but not genotypes. Simulations are initialized with each deme initially maximally fit (i.e., all deleterious allele frequencies are zero). In each generation, allele frequencies and population sizes are updated to reflect the effects of migration, mutation, selection and reproduction, as described in Online Supplement A. The metapopulation is allowed to equilibrate and the equilibrium load and population size are computed by averaging over multiple time points and simulation replicates.

### Choice of parameters

To guide the choice of parameters, we look to broad estimates across a range of organisms. For example, the genome-wide rate of deleterious mutations per individual per generation is estimated to be 0.1−1 in multicellular eukaryotes (Lynch et al., 1999), 1.2−1.4 in *Drosophila* (Haag-Liautard et al., 2007), 0.25−2.5 for *C. elegans* (Denver et al., 2004) and 2.2 for humans (Keightley, 2012), the latter being an unusually high value. Estimates of fitness effects of new mutations are similarly varied: while some experimental studies have shown that a sizeable fraction (*<*15%) of mutations are likely to be lethal (Eyre-Walker et al., 2006; Mukai et al., 1972), there is consensus that small effect mutations (with effect ≲5% on quantitative traits) have higher densities (Lyman et al., 1996; Lynch et al., 1999; Mukai, 1964; Mukai et al., 1972). Dominance coefficients are harder to estimate for weakly deleterious mutations but may be ~0.2 for moderately deleterious variants (Charlesworth and Charlesworth, 1987). In addition, several studies point to a negative relationship between *s* and *h*, with mutations of small effect being almost additive and those of large effect being almost recessive (Lynch et al., 1999; Simmons and Crow, 1977).

Finally, indirect measures (e.g, *F*_*ST*_) have reported moderate to high levels of gene flow in plants and mammals with typical values between closely related species being 0.05−0.2, which corresponds roughly to between 1−5 migrants between local populations per generation. It is important to note that in populations that are on the verge of extinction, gene flow may be much less than this (Casas-Marce et al., 2013; Frankham et al., 2002; Szczecińska et al., 2016) and indeed one of the goals of this study is to identify critical levels of migration required to prevent population collapse. Thus, most of the study will focus on low to moderate levels of gene flow.

The most difficult parameter to estimate is the baseline growth rate, *r*_0_ (see also Discussion). This enters into the composite parameter 2*L*(*u/r*_0_), which is a measure of the ‘hardness’ of selection. We restrict ourselves to values less than 1, as extinction is guaranteed for 2*L*(*u/r*_0_)*>*1 (regardless of gene flow).

## Results

### Soft selection

We first consider soft selection, where each deme is at carrying capacity *K*, regardless of mean fitness. If loci evolve independently, then the equilibrium allele frequency distribution 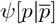 at any locus in any deme depends only on the scaled parameters *Ks, Ku, Km* and the dominance coefficient *h*, and is given by eq. (3), conditional on 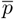, the mean allele frequency across all demes. As discussed above, 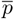 can be obtained self-consistently by numerically solving 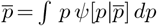.

The soft selection model has been analysed in earlier work assuming either very weak or strong selection, i.e., *Ks*≪1 (Whitlock, 2002), or *Ksh*+*Km* at least 5 (Glémin et al., 2003; Roze, 2015). These limits are useful to consider as they yield simpler intuition and allow for explicit analytical results: for instance, when selection is much weaker than local drift (i.e., *Ks*≪1), probabilities of identity by descent of two or more genes are approximately the same as under the *neutral* island model, allowing one to approximate 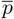 in terms of (essentially) neutral *F*_*ST*_ (Whitlock, 2002). In this limit, selection is only efficient at the level of the population as a whole and not within demes, some of which may be locally fixed for the deleterious allele. By contrast, for large *Ksh*, deleterious alleles segregate at low frequencies within any deme and the allele frequency distribution is concentrated about 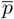. Now, gene flow only further narrows the distribution of allele frequencies by further increasing the effective size of local demes and the efficiency of selection (Glémin et al., 2003; Roze, 2015). Here, we focus on *moderately* deleterious alleles with *Ks*~1 (which contribute the most to drift load in an isolated population). As we argue in Online Supplement B, the effects of selection in this regime are more subtle and not captured by either intuitive description above.

Beyond these more conceptual issues, the concrete question we address is: how much gene flow between subpopulations is required to alleviate drift load at a single locus under soft selection, and how does this depend on the (scaled) selection strength *Ks* and dominance coefficient *h* at the locus? We then consider how load per locus is affected by multilocus associations between many *unlinked* deleterious alleles.

To begin with, consider the case of isolated demes (*Km*=0). For *Ku*≪1, any deme will be close to fixation for one or other allele at any locus, allowing us to use a ‘fixed state’ approximation (Barton and Olusanya, 2022; Szép et al., 2021). This approximates the U-shaped allele frequency distribution *ψ*(*p*) by the sum of two delta functions (spikes)– one at *p*=0 (corresponding to near-fixation of the wildtype allele) and the other at *p*=1 (near-fixation of deleterious allele). The relative weights of these two spikes are: 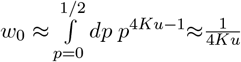, and 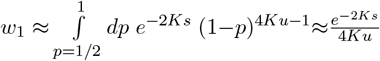, regardless of *h*. Thus, the expected deleterious allele frequency is 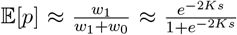, and the expected load *G* (scaled by the deterministic expectation 2*u*) is: 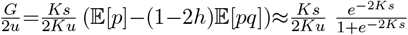, assuming 𝔼 [*pq*] ≈ 0. It follows that the maximum contribution to load is from loci with *Ks*≈0.64, independent of *h*, with load per locus being 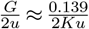 for this value of *Ks*. In other words, load due to moderately selected loci (say, *Ks*~ 0.64) would be ~70 (or ~700) times the deterministic expectation 2*u* if *Ku*=0.001 (or *Ku*=0.0001), which corresponds to *u/s*=0.0015 (or *u/s*=0.00015). Thus, drift can inflate load due to moderately selected loci by a factor of several hundreds or thousands in isolated populations (see also Kondrashov, 1995).

Now consider the effect of gene flow between subpopulations: fig. 1a shows load per locus *G* (scaled by 2*u*) and the expected deleterious allele frequency (inset) as a function of *Km*, for *Ks*=0.64 and various values of *h*. Symbols depict predictions of the full diffusion framework (eq. (3)), and lines those of a simpler ‘moderate selection’ approximation, details of which are outlined in Online Supplement B.

**Figure 1.**
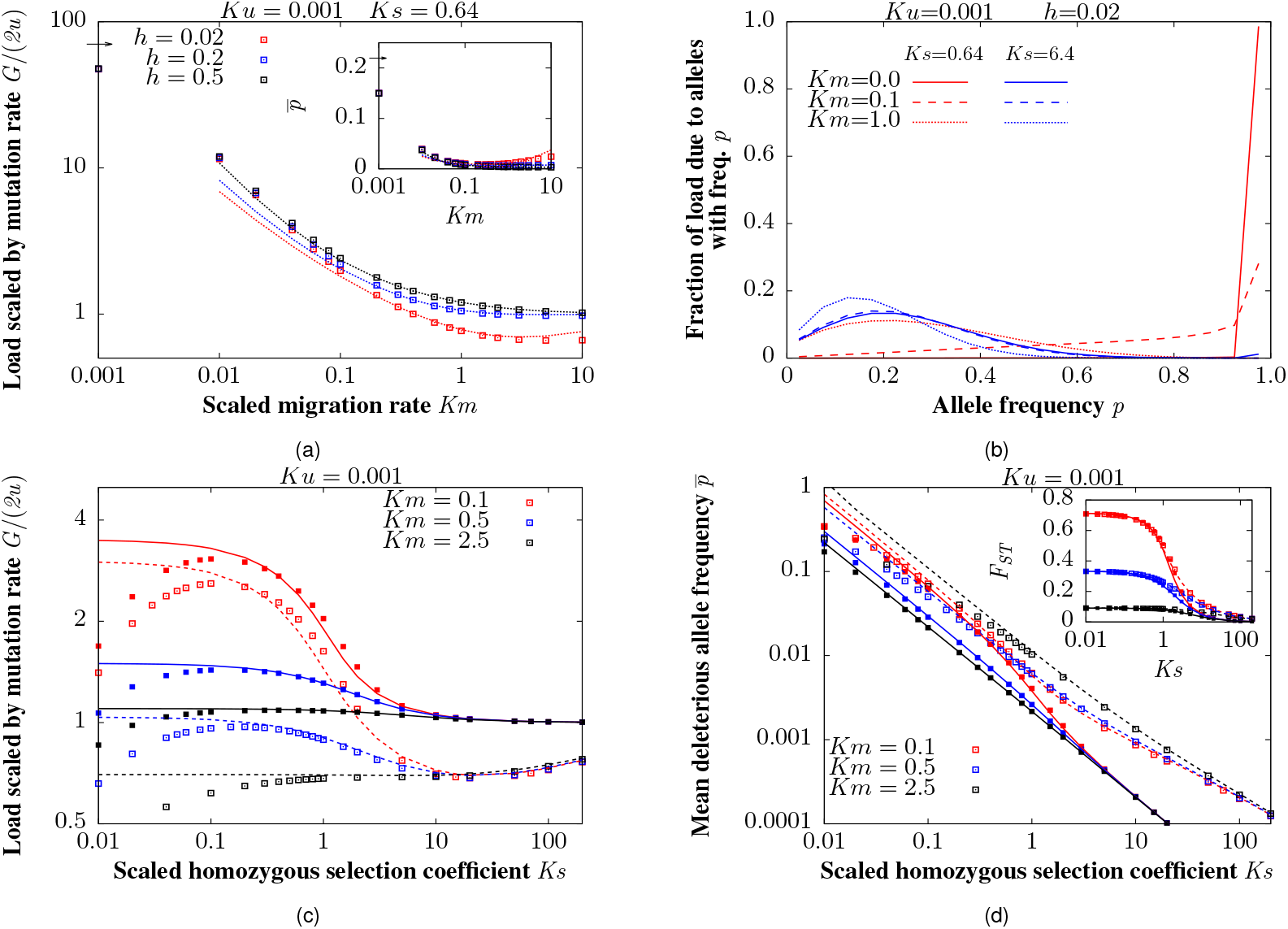
Mutation-selection-drift-migration equilibrium at a single locus under the infinite-island model with soft selection. (a) Main plot and inset show respectively the expected per locus load *G* (scaled by 2*u*) and the mean deleterious allele frequency 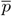 vs. *Km*, the number of migrants per deme per generation, for various values of *h*, for *Ks*=0.64 (which is the *Ks* value for which load is maximum in an isolated population). Symbols depict results of the diffusion approximation (eq. (3)), while lines represent predictions of the ‘moderate selection’ approximation (eq. S7a, Online Supplement B). Horizontal arrows on the y-axis represent the corresponding diffusion predictions for *Km*=0. (b) The fraction of total load that is due to alleles with frequency in the interval between *p* and *p*+Δ*p* (with Δ*p*=0.05) for various values of *Km* (solid, dashed and dotted lines), for weakly selected (*Ks*=0.64; red) and strongly selected (*Ks*=6.4; blue) loci. Predictions are based on the diffusion approximation, i.e., obtained by integrating over the allele frequency distribution in eq. (3). (c) Expected per locus load *G* (scaled by 2*u*) vs. *Ks*, the homozygous selective effect scaled by drift, for various values of *Km* (different colours) for *h*=0.5 (filled symbols) and (open symbols). (d) Mean deleterious allele frequency 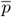 (main plot) and the expected *F*_*ST*_ at selected loci (inset) vs. *Ks*, for various *Km* (different colours) for *h*=0.5 (filled symbols) and *h*=0.02 (open symbols). In both (c) and (d), symbols depict results of the diffusion approximation (eq. (3)) and lines show the ‘moderate selection’ predictions. All plots are with *Ku*=0.001.

**Figure 2.**
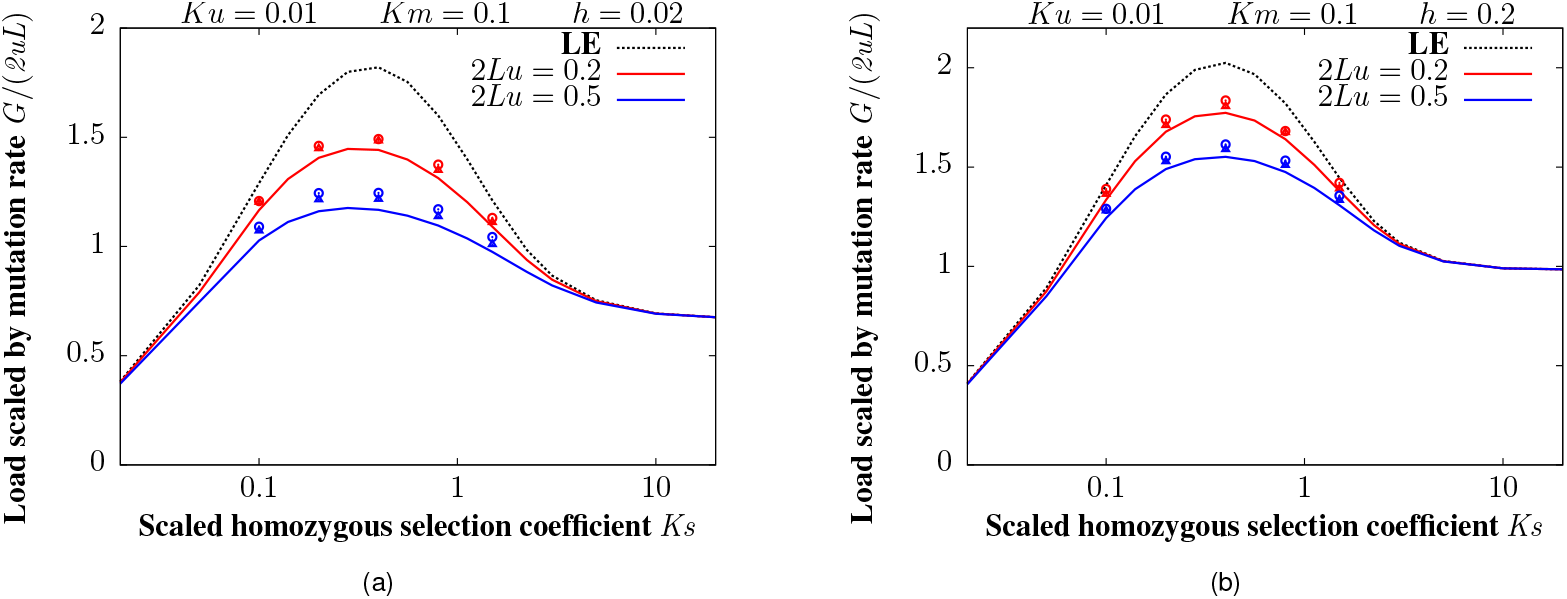
E ffect of multi-locus heterosis on load. E quilibrium load (scaled by total mutation rate 2*Lu*) vs. *Ks*, in a population with deleterious mutations occurring at *L* biallelic equal-effect loci with (a) *h*=0.02 and (b) *h*=0.2, for *Km*=0.1. Symbols show results of individual-based simulations for two values of total mutation rate 2*Lu*, and two values of *L* for each 2*Lu*. Red circles and triangles depict respectively *L*=1000, *u*=10^−4^ and *L*=2000, *u*=5 *×* 10^−5^ both of which correspond to 2*Lu*=0.2; blue circles and triangles depict *L*=2500, *u*=10^−4^ and *L*=5000, *u*=5 *×* ^−5^, which correspond to 2*Lu*=0.5. For each value of *L* and *u*, the carrying capacity *K* and migration rate *m* are chosen such that *Ku*=0.01 and *Km*=0.1. For a given *K*, the scaled selection coefficient *Ks* is varied by varying *s*. Dashed curves depict single-locus predictions (obtained using eq. (3)). Solid curves show predictions that account for multi-locus heterosis using effective migration rates; these predictions are independent of *L* for a fixed *Lu, u/s, Ks* and *Km*. Solid and dashed curves differ markedly, i.e., multi-locus heterosis causes a significant reduction in load for intermediate *Ks*, small *h* and large *Lu*.

**Figure 3.**
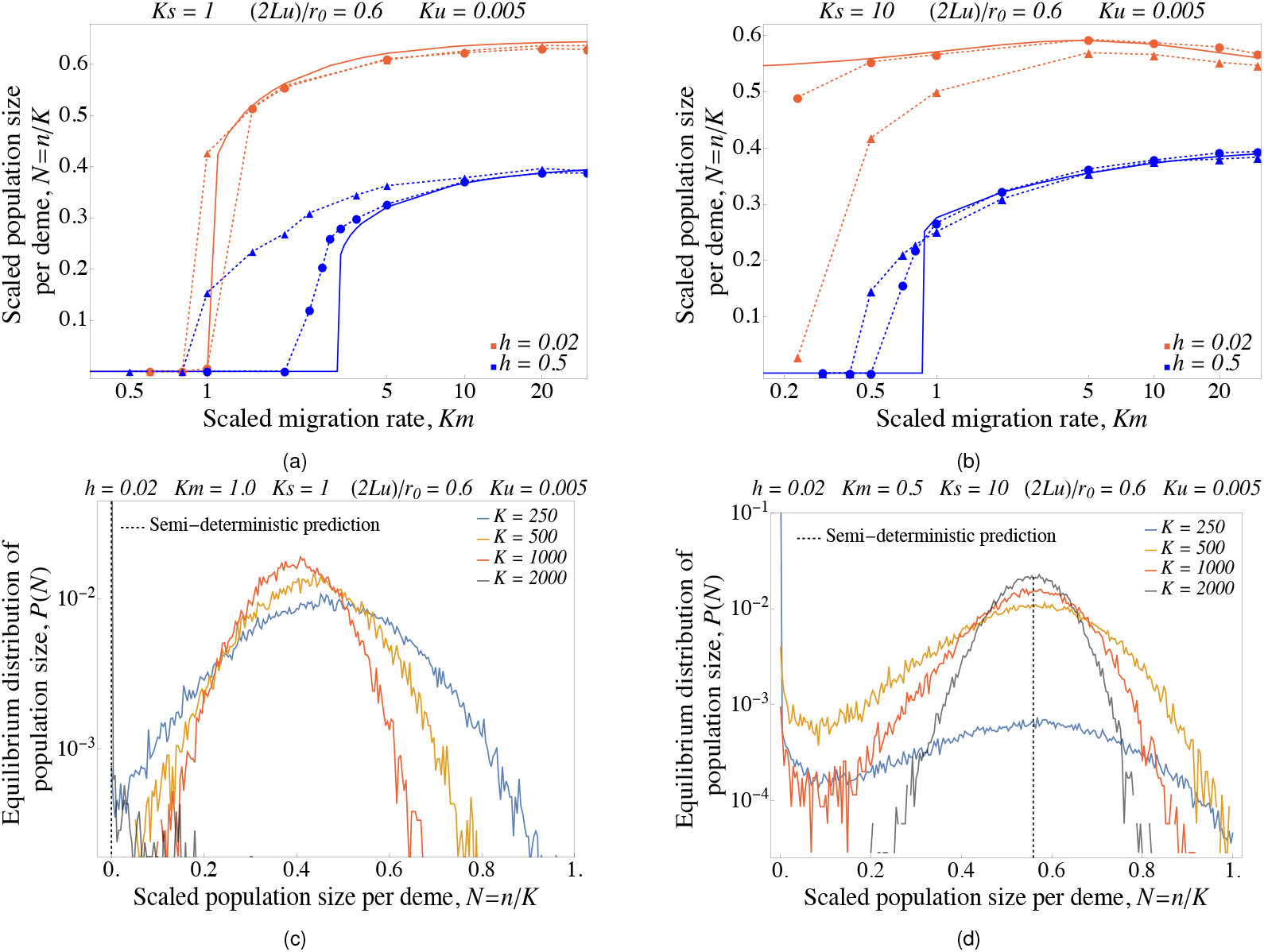
Comparison of simulation results with semi-deterministic predictions. Mean (scaled) population size *N* per deme plotted against *Km* for (a) *Ks*=1 and (b) *Ks*=10 for nearly recessive (*h*=0.02) and additive (*h*=0.5) alleles; other scaled parameters are: *Ku*=0.005 and (2*Lu*)*/r*_0_=0.6. Solid lines show predictions of the semi-deterministic approximation; symbols connected by dashed lines show results of allele frequency simulations with carrying capacity per deme *K* either 500 (triangles) or 3000 (circles). (c)-(d) Equilibrium distribution *P* (*N*) of population sizes *N*, as obtained from simulations, for various values of *K*, for two different scaled parameter combinations for which the semi-deterministic approximation predicts extinction (in (c)) vs. stable populations (in (d)). The scaled parameters *Ks, Ku, Km*, (2*Lu*)*/r*_0_) are held constant as *K* is varied (by simultaneously varying *s, u, L*, and *m*). The scaled parameters have values: (c) *Ks*=1, *Km*=1, or (d) *Ks*=10, *Km*=0.5, with other parameters being *h*=0.02, *Ku*=0.005 and (2*Lu*)*/r*_0_=0.6, as above. Black vertical dashed lines represent the semi-deterministic prediction for *N* : this is *N* =0 in (c) and *N* =0.56 in (d). The distribution *P* (*N*) becomes sharply peaked around the semi-deterministic prediction as *K* increases in both (c) and (d). All simulations were performed with 100 demes and *r*_0_=0.1.

Figure 1a shows that a very low level of gene flow (*Km*~0.1) is enough to dramatically reduce drift load, regardless of dominance. For instance, *G/*(2*u*) falls from ~70 at *Km*=0 to 4−5 at *Km*=0.1 for both nearly recessive (*h*=0.02) and co-dominant (*h*=0.5) deleterious alleles, for *Ku*=0.001. In both cases, the expected allele frequency 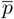 declines from 0.22 at *Km*=0 to ~0.01 at *Km*=0.1 (inset), while heterozygosity hardly increases (from ~0.002 to 0.003−0.004). Thus the reduction in load at low levels of migration is almost entirely due to the decline in the number of fixed deleterious alleles, with little to no change in the number of segregating alleles.

Higher levels of migration further reduce load, though in the case of recessive alleles, this is partially offset by an increase in heterozygosity (which results in less efficient purging). Thus, for recessive alleles, load is minimum at *Km*≈4 and rises again as *Km* increases further, approaching the level expected in a panmictic population at large *Km*. For the parameters shown in fig. 1a, the reduction in load due to purging at intermediate *Km* is rather modest and hardly visible on the scale of the plot; purging has a more substantial effect when *u/*(*hs*) and *Ks* are smaller, and provided *h<*1*/*4 (Whitlock, 2002).

Increased gene flow also shifts the frequency spectrum of alleles that contribute to load – for *Ks*≲0.1, the predominant contribution at *Km*=0.1 is from fixed or nearly fixed alleles. However, one migrant per deme per generation (*Km*=1) is already enough to prevent fixation of deleterious alleles, so that load is entirely due to alleles segregating at intermediate frequencies at this higher migration level (dashed vs. dotted red curves in fig. 1b). As expected, gene flow has a much weaker effect at strongly selected loci: for *Ks*=6.4, load is entirely due to segregating alleles regardless of *Km* (blue curves). Thus, gene flow only causes a minor reduction in load at strongly selected loci (fig. 1c), and may even be detrimental if it hinders purging (in the case of recessive deleterious alleles).

Moreover, even low levels of gene flow tend to ‘even out’ the contributions of alleles with different selection coefficients to load, so that moderately deleterious alleles no longer contribute disproportionately (as in isolated populations). For instance, with *Km*=0.1, load is maximum for *Ks* ≈ 0.1 (regardless of *h*), but is only about ~3 times larger than the deterministic expectation (fig. 1c). By contrast, in isolated populations, it may be several hundred or thousand times larger (see above). With modest levels of gene flow (*Km*=0.5), the increase in load for intermediate *Ks* is weaker still, while ≳2 migrants per generation are enough for load to be more or less independent of *Ks*, as in a large, panmictic population.

One can also ask: are the processes underlying this dramatic reduction in load (even with low levels of gene flow) qualitatively different for alleles with different selective effects? Expressing load as 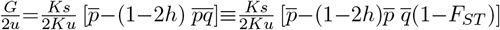, it follows that for recessive alleles (*h<*1*/*2), load declines if the mean deleterious allele frequency 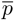 and/or *F*_*ST*_ at the selected locus decrease. A decline in *F*_*ST*_ (for a fixed 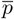), in turn, reflects a change in the allele frequency distribution from a more *U*-shaped distribution (wherein a proportion 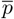 of demes are nearly fixed for the deleterious allele) to a unimodal distribution (wherein the deleterious allele segregates at a low frequency close to 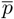 in all demes).

Figure 1d shows 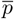 and *F*_*ST*_ vs. *Ks* (main plot and inset), for various values of *Km*, for co-dominant and nearly recessive alleles (filled vs. open symbols). Note that while selection as weak as *Ks*≳0.05 is already quite efficient at reducing 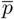, the mean number of deleterious alleles (say, under low gene flow, *Km*=0.1), it has little effect on *F*_*ST*_ (i.e., on the probability of local fixation within demes, given 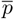) unless *Ks*≳0.5. For example, *F*_*ST*_ is reduced by only about 15% with respect to its neutral value for alleles with *Ks*=0.5, but by *>*50% for *Ks*=2, for both values of *h*. Thus, selection for increased heterozygosity within demes (as reflected in a decrease in *F*_*ST*_) does not markedly influence the evolutionary dynamics of alleles with *Ks*≲1 (which suffer the most severe inflation of load in isolated populations) but can be important for moderately deleterious alleles with 1≲*Ks*≲5 (which may still contribute substantially to load).

In Online Supplement B, we introduce a ‘moderate selection’ approximation, which accounts for the effects of selection on *F*_*ST*_, but assumes that (appropriately scaled) higher cumulants of the allele frequency distribution are related to *F*_*ST*_ in the same way as under the neutral island model. The predictions of this approximation are shown by lines in fig. 1a, 1c and 1d, and appear to match the full diffusion (symbols) quite well for 0.5≲*Ks*≲5. This suggests that moderate selection essentially changes pairwise coalescence times (to different extents within and between demes), without appreciably affecting other statistical properties of the genealogy, e.g., the degree to which branching is skewed or asymmetric, at loci subject to purifying selection (see also Online Supplement B).

### Effect of multilocus associations on load

So far, we have neglected statistical associations, i.e., linkage and identity disequilibrium between deleterious alleles at different loci. These may be substantial if, for instance, different subpopulations are nearly fixed for partially recessive deleterious alleles at many different loci, so that *F* 1 individuals (offspring of migrants and residents) have higher fitness than either parent, giving rise to heterosis. In such a scenario, the extent of gene flow at any locus depends not only on *Km*, the average number of migrants between demes, but also on the relative fitness of migrants and their descendants. The fitness of these descendants may be significantly higher than that of residents if there is high differentiation between subpopulations (i.e., if migration is low or *F*_*ST*_ high), if deleterious alleles segregate at very many loci (2*Lu* is large and *Ks* small) and are mostly recessive (*h* is small).

More specifically, heterosis causes immigrant alleles to be transmitted to the next generations at a higher rate than alleles carried by residents, resulting in an *effective* migration rate that is higher than the raw migration rate (see also Ingvarsson and Whitlock (2000)), which reduces per-locus load. For a genome with *L* unlinked loci with equal selective effects *s* and dominance coefficients *h*, and assuming *s*≪1, the effective migration rate at any locus is shown by Zwaenepoel et al. (2023) to be approximately:

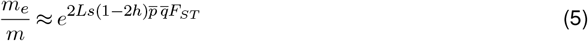

Following Sachdeva (2022) and Zwaenepoel et al. (2023), we approximate the effects of multi-locus heterosis on allele frequencies by assuming that these follow the equilibrium distribution in eq. (3), but with the raw migration rate replaced by an effective migration rate which itself depends on the expected allele frequency 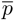 and *F*_*ST*_ (or equivalently, 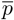 and E [*pq*]) through eq. (5). This allows us to numerically solve for 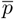 and *F*_*ST*_, yielding the theoretical predictions (solid lines) in fig. 2.

Figure 2 shows total load scaled by 2*Lu* or equivalently, load per locus scaled by 2*u*, as a function of *Ks* for low migration (*Km*=0.1), for two values of the total mutation rate (2*Lu*=0.2 in red and 2*Lu*=0.5 in blue) and two dominance coefficients (*h*=0.02 in fig. 2a and *h*=0.2 in fig. 2b). For each value of 2*Lu*, we further simulate two values of *L*, scaling down *s, u* and *m*, and scaling up *K* as we increase *L*, so that *Ks, Km, Ku* and *Ls* remain constant. Symbols depict results of individual-based simulations for a metapopulation with 100 demes; dashed curves show single-locus predictions (obtained using eq. (3)) that do not account for multilocus heterosis; solid curves represent predictions that account for heterosis using effective migration rates: note that these depend on *L* only via the combination *Ls* (or alternatively, *Lu* for a given *u/s*). As expected, there is better agreement between simulations and theory for smaller values of *s* (or alternatively, larger *L*), for a given total mutation rate, as the expression for effective migration rate in eq. (5) becomes more accurate as *s*→0. As can be seen in fig. 2, load per locus is most strongly reduced by multilocus heterosis when 2*Lu* is large, and for loci with small *h* and intermediate *Ks*. Moreover, the effects of multilocus heterosis become weaker with increasing migration which reduces *F*_*ST*_, i.e., differentiation between demes, and consequently the extent of heterosis (see also Roze, 2015).

Note that the predictions based on effective migration rates that we describe above (and depict in Fig. fig. 2) account for the effects of fitness variation between demes at any focal locus but not those of fitness variance within demes. The latter is known to reduce the effective population size, thus, further impairing the efficiency of selection and generating Hill-Robertson interference between deleterious alleles (McVean and Charlesworth, 2000). However, for unlinked loci, this effect is rather modest– for instance, for *Ns*≫1, the effective population size is ~ *e*^−8*Lu hs*^ times the true population size (Charlesworth, 2012), which implies a 1% reduction if, say, 2*Lu*=0.5, *s*=0.01 and *h*=1*/*2.

### Hard selection

The analysis of soft selection in the previous section bears out ‘one migrant per generation’ as an approximate threshold beyond which drift load (due to random fixation of deleterious alleles) ceases to be significant (see e.g., fig. 1a). This is true regardless of *h* and even for moderately deleterious alleles that would otherwise contribute disproportionately to load. Here, we ask: what is the analogous criterion under hard selection, where feedback between population size and load can lead to extinction.

Under soft selection, the efficiency of selection is governed by *Ks*, selection per locus relative to local drift. However, under hard selection, population sizes may be less than *K*, so that (scaled) selection per locus is *ns*=*NKs* (where *n* and *N* =*n/K* are the unscaled and scaled population size of a deme respectively). In general, we expect 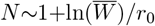 (eq. (4)), i.e., population sizes to be reduced relative to carrying capacity by an amount equal to the total load − 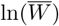 scaled by the baseline growth rate *r*_0_. In the absence of drift, − 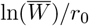 is simply (2*Lu*)*/r*_0_ (which we use as a measure of the ‘hardness’ of selection, i.e., of the strength of the eco-evo feedback). However, drift will typically increase load (unless alleles are highly recessive) and thus depress population size. This results in a positive feedback between increase in load per locus (which is governed by per locus effects *NKs* and *h*) and a decrease in population size (which is governed by total load due to all *L* loci). Thus, the strength of the positive eco-evo feedback is mediated by the genetic architecture of load, i.e., the numbers, effect sizes and dominance coefficients of deleterious alleles. In the following, we will focus on the case of equal effect loci; scenarios involving a distribution of fitness effects are explored in Online Supplement C.

The analysis in this section is based on the semi-deterministic approximation. We first test its accuracy by comparing against allele frequency (linkage equilibrium) simulations (fig. 3) and then use the semi-deterministic analysis to identify critical migration thresholds below which the metapopulation becomes extinct, i.e., *N* =*n/K* goes to zero. We explore how these thresholds depend on *Ks, h* and (2*Lu*)*/r*_0_ (fig. 4). Finally, we consider parameter regimes for which the metapopulation is stable (i.e., not extinct) and explore how equilibrium load and population size depend on various parameters (fig. 5).

**Figure 4.**
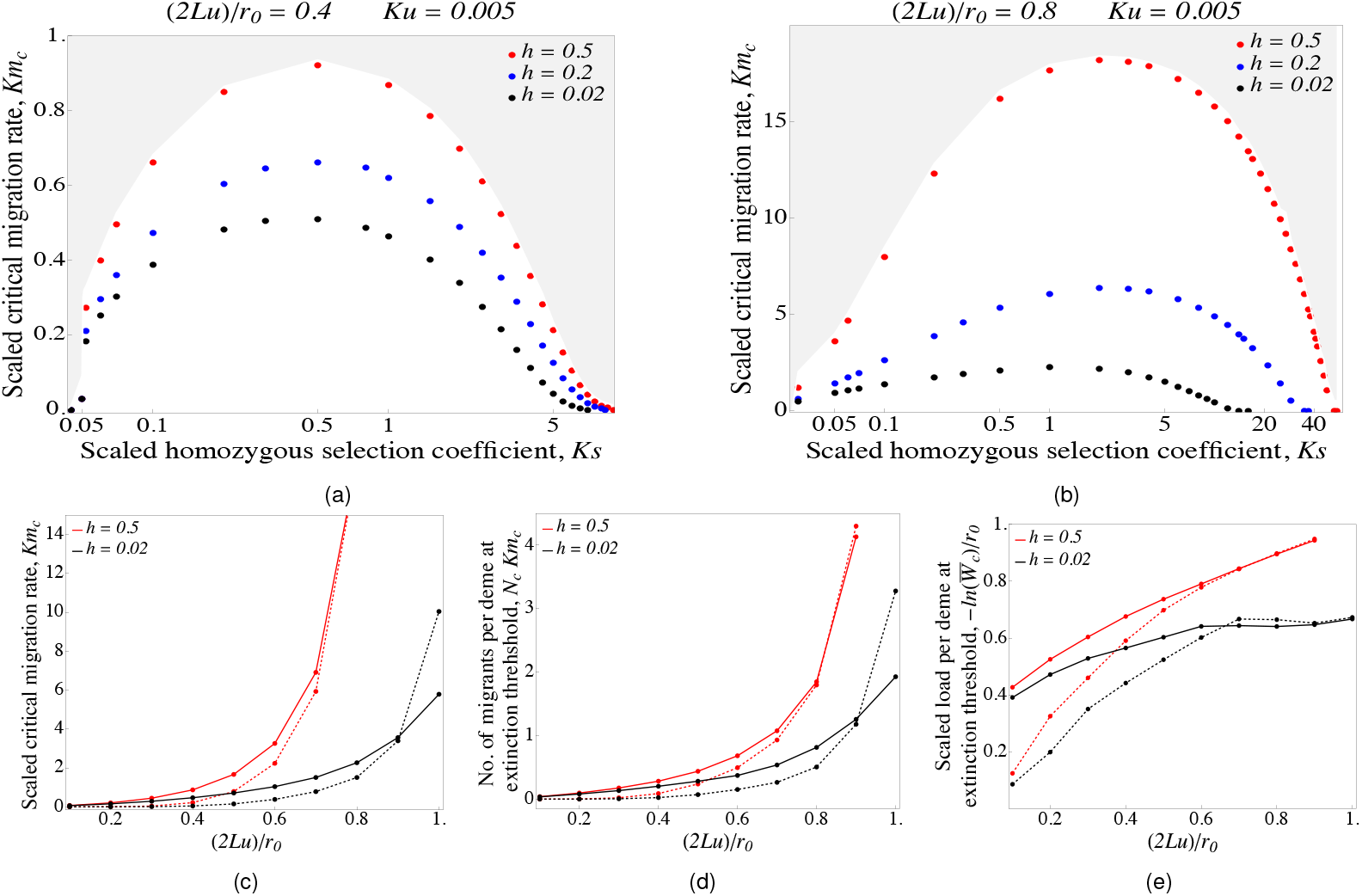
(a)-(b) Critical migration thresholds *Km*_*c*_ (below which the metapopulation goes extinct) as a function of *Ks* for (a) (2*Lu*)*/r*_0_=0.4 (corresponding to moderately hard selection) and (b) (2*Lu*)*/r*_0_=0.8 (much harder selection), for *Ku*=0.005. Figures (c)-(e) show: (c) the critical migration threshold, *Km*_*c*_ (d) the number of migrants per deme, *N*_*c*_*Km*_*c*_, just above the critical migration threshold, and (e) the scaled load per deme, − ln 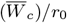,, just above the critical migration threshold, as a function of (2*Lu*)*/r*_0_, which is a measure of the hardness of selection. Solid and dashed lines represent results with *Ks*=1 and *Ks*=5 respectively; different colors correspond to different values of *h*. All predictions are obtained from the semi-deterministic analysis (using eq. (4)), which is accurate when carrying capacity per deme is large.

**Figure 5.**
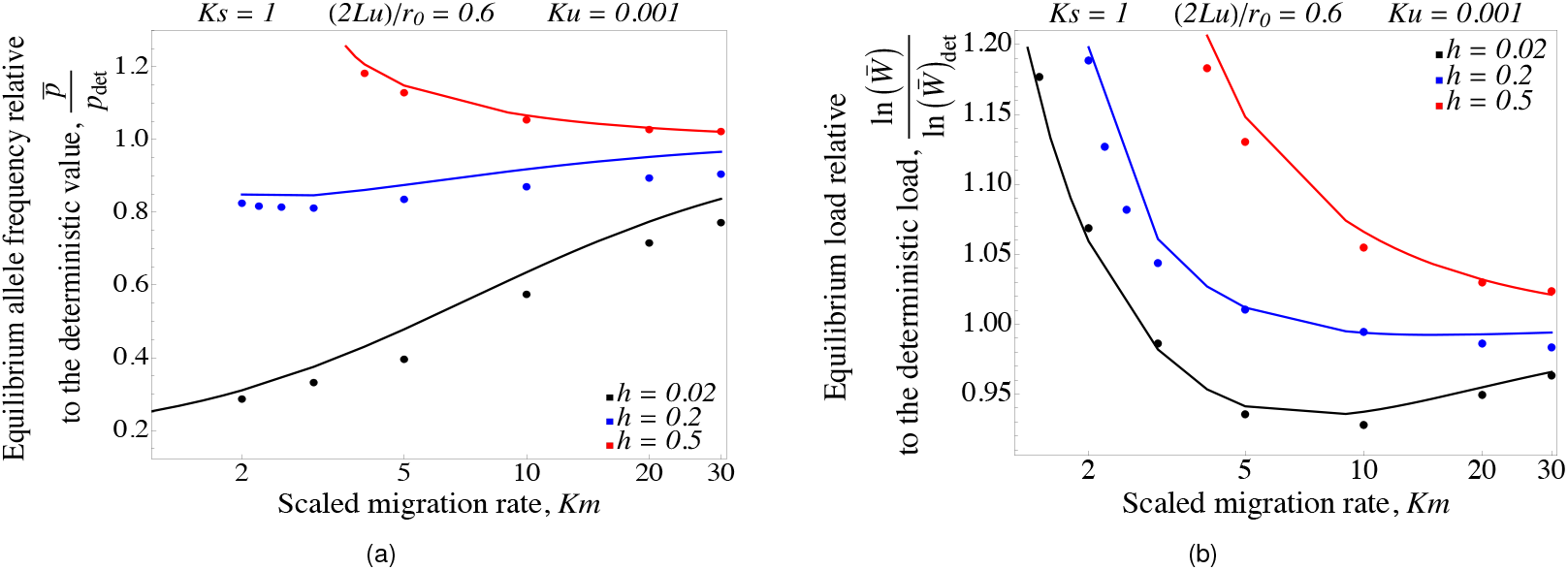
(a) E quilibrium frequency 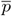 and (b) equilibrium load − ln 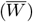 (relative to *p*_*det*_ and ln 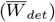, the deterministic allele frequency and load in a large undivided population), plotted against *Km*, for *Km>Km*_*c*_. Filled circles show simulation results (using allele frequency simulations) and solid lines show results from the semi-deterministic approximation. The scaled parameters are *Ku*=0.001, *Ks*=1 and (2*Lu*)*/r*_0_=0.6; simulations are run with 100 demes with *K*=3000 and *r*_0_=0.1.

### Critical migration thresholds for metapopulation extinction

Figure 3a shows the mean population size per deme as a function of *Km*, the average number of migrants per deme (for demes at carrying capacity). Deleterious mutations of equal effect occur at *L* loci with (2*Lu*)*/r*_0_=0.6; thus selection is fairly hard. We compare scenarios where deleterious mutations have moderate (*Ks*=1) vs. strong (*Ks*=10) effects (fig. 3a vs. 3b), and also recessive vs. additive effects (red vs. blue) for each *Ks*. The solid lines show semi-deterministic predictions (derived using eq. (4)) and symbols connected by dashed lines show simulation results. In order to understand the effects of demographic stochasticity, we run simulations for various values of *r*_0_*K* (which is varied by changing *K*, with *r*_0_ fixed at 0.1), while keeping all scaled parameters (*Ks, Ku, Km* and (2*Lu*)*/r*_0_) fixed. This is done by simultaneously reducing *s, m*, and *u* and increasing *L*, as *K* is increased. These simulations establish that (as expected) the semi-deterministic approximation becomes increasingly accurate in the limit *r*_0_*K*→∞, i.e., as demographic fluctuations become weaker (compare triangles vs. circles which correspond to *K*=500 and *K*=3000 respectively).

For three out of the four parameter combinations in figs. 3a and 3b, we observe a critical migration threshold (denoted by *Km*_*c*_) below which the metapopulation collapses. For *Km<Km*_*c*_, patches accumulate deleterious alleles and dwindle in size faster than they can be rescued by migration, leading to their extinction. A noticeable feature of figs. 3a and 3b is the discrepancy between simulations and the semi-deterministic predictions, especially for *Km*_*c*_. However, as above, this discrepancy is lower for larger *K*. Note also that for large *K*, i.e., when the semi-deterministic approximation is valid, extinction thresholds resemble ‘tipping points’, where a slight drop in migration sets in motion a positive feedback between increasing load and decreasing population size which causes relatively large populations to collapse abruptly.

The critical threshold *Km*_*c*_ is larger for smaller values of *Ks* and larger *h*. In other words, higher levels of migration are required to sustain populations when selection per locus is weak relative to drift and if alleles are additive rather than recessive (since purging is more effective for recessive alleles). On the flip side, when alleles are recessive, the population size per deme is higher and load correspondingly lower at the critical threshold *Km*_*c*_. For example, in Figure 3a, extinction already ensues for 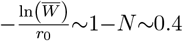 if load is due to recessive alleles, since even a slight increase in homozygosity (due to reduced migration and/or declining population sizes) causes load to increase substantially, resulting in stronger eco-evo feedback. By contrast, populations can sustain much higher levels of load *without* collapsing, when load is due to additive alleles (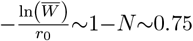 at *Km*_*c*_; blue lines in figs. 3a and 3b).

Interestingly, the critical migration threshold *Km*_*c*_ in simulations with smaller *K* is *lower* than the semi-deterministic prediction (which assumes *r*_0_*K*≫1), all other scaled parameters being the same. In other words, given *Ks*, (2*Lu*)*/r*_0_, *Ku*, smaller patches with lower carrying capacities *K* (and stronger demographic fluctuations) require fewer migrants per generation to escape extinction. A related finding is that just above the extinction threshold, scaled population sizes *N* =*n/K* are actually higher when *K* is smaller. These observations appear somewhat puzzling, as one might expect demographic stochasticity to promote extinction of individual patches. However, this is counterbalanced by the fact that stronger demographic fluctuations result in a broader distribution of deme sizes overall. This allows some demes within the metapopulation to function as demographic sources— migration from these can then rescue other demes with declining populations. Moreover, the demographic, i.e., number-boosting effects of migration (as captured by the second term in eq. (2b)) are also stronger for smaller *K* (for a fixed *Km*). More specifically, if *K* is small, migration can rescue individual patches more effectively not just by preventing fixation of deleterious alleles (i.e., via a genetic effect) but also by directly increasing population numbers (i.e., via a demographic effect), where increased population numbers further hinder deleterious allele accumulation (Sachdeva et al., 2022).

These effects are illustrated in fig. 3c, which shows the equilibrium distribution of deme sizes for different values of *K*, for *Ks*=1, *h*=0.02, and *Km*=1 (which is just below the semi-deterministic prediction for *Km*_*c*_). When carrying capacities are small (say *K*=250; blue), the distribution of scaled population sizes is rather broad. For larger *K* (i.e., with weaker demographic fluctuations), the distribution becomes more narrow, with fewer exceptionally large patches that can potentially act as demographic sources. As a result, the entire distribution *shifts* towards smaller *N*. At still larger values of *K* and, consequently, with narrower distributions, the entire population collapses, yielding an equilibrium size *N* =0 (which matches the semi-deterministic prediction).

The effect of demographic fluctuations is, however, qualitatively different in parameter regimes where the semi-deterministic approximation predicts viable populations, i.e., non-zero equilibrium size (Figure 3d). At low *K* (e.g. *K*=250; blue in Figure 3d), population sizes show a bimodal distribution, with a small fraction of demes close to extinction and the remaining fraction peaked at a non-zero *N*_*_. Now, as *K* increases, the distribution around *N*_*_ becomes narrower and smaller sub-populations become less prone to collapse, resulting in fewer nearly extinct demes. In fact, for *K*=2000, all demes have sizes close to *N*_*_ ~ 0.56 which coincides with the semi-deterministic prediction (dashed vertical black line). Similar plots are shown for *h*=0.5 in Online Supplement D.

We now investigate how *Km*_*c*_, the critical level of migration required to prevent population collapse, depends on the architecture of load, i.e., the (scaled) selection and dominance coefficients per locus and the total deleterious mutation rate relative to baseline growth rate. We only show predictions of the semi-deterministic approximation, which is expected to be increasingly accurate for large *r*_0_*K*. Importantly, we always start from a state in which the metapopulation is large and well-adapted. An alternative scenario (not considered here) is one where the metapopulation grows (or fails to grow) starting from a few individuals per patch. In this case, much higher migration between patches is required to alleviate so-called genetic Allee effects (Courchamp et al., 2008; Fauvergue et al., 2012), wherein populations smaller than a critical size are unable to initiate growth due to high inbreeding load (see also Sachdeva et al., 2022).

Figures 4a and 4b show that there is a non-monotonic relationship between the critical migration threshold *m*_*c*_ and selection per locus. Moreover, *m*_*c*_ is higher for larger values of *h*. This reflects the fact that load per locus is largest for alleles of moderate selective effect *Ks*, and for additive as opposed to recessive alleles (fig. 1c); thus the level of migration required to alleviate load and prevent extinction is also highest for intermediate values of *Ks* and larger *h*. Interestingly, the critical load threshold, 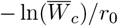, in populations on the brink of collapse (i.e., just above *Km*_*c*_) is lower when *Ks* is larger (dashed vs. solid lines in fig. 4e). In other words, the maximum amount of load (relative to baseline growth rate) that populations can sustain without collapsing is lower if selection per locus is stronger. This is because while strongly deleterious alleles are less likely to fix, the loss in fitness when fixation does occur is higher (and the resultant eco-evo feedback between load and population size stronger) for alleles with larger *s*.

A key determinant of the eco-evo feedback is (2*Lu*)*/r*_0_, the total deleterious mutation rate (which determines load in a large population) relative to *r*_0_. As discussed above, (2*Lu*)*/r*_0_ can be viewed as a proxy for the ‘hardness’ of selection. Figures 4a and 4b depict *Km*_*c*_ for (2*Lu*)*/r*_0_=0.4 and (2*Lu*)*/r*_0_=0.8 respectively, and show that critical migration thresholds required to prevent collapse can increase dramatically as selection becomes more hard. Moreover, the relatively small difference in load per locus between additive and recessive alleles that we observe at a given level of migration under soft selection (fig. 1c) can translate into rather different critical migration thresholds for the two kinds of alleles under hard selection. This is especially marked when (2*Lu*)*/r*_0_ is higher (fig. 4b), as a small increase in fixation probability and load per locus can translate into a substantial change in total load and population size, which further increases fixation probabilities, resulting in very strong feedback effects.

Some care is needed in interpreting *Km*_*c*_: this is best thought of as the minimum rate of migration (scaled by *K*) that allows the metapopulation to avoid collapse. It is thus equal to the number of migrants per deme that one would observe in a *hypothetical* metapopulation at carrying capacity, given this migration rate. It is not the number of migrants per deme in the *actual* metapopulation just above the critical migration threshold, which is given by *N*_*c*_*Km*_*c*_, where *N*_*c*_ is the (scaled) population size per deme at this migration threshold. Figures 4d and 4e show the average number of migrants *N*_*c*_*Km*_*c*_ and the average (scaled) load per deme, − ln 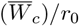, at the critical extinction threshold, as a function of (2*Lu*)*/r*_0_, for the various combinations of *Ks* and *h*. Thus, for instance, for the parameters (2*Lu*)*/r*_0_=0.6, *Ks*=1 and *h*=0.5, a minimum of *N*_*c*_*Km*_*c*_ ~ 0.75 migrants per deme per generation are required for the metapopulation to avoid extinction (fig. 4d). The load per deme in this metapopulation just above the extinction threshold is − ln 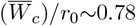 (fig. 4e), and the (scaled) population size per deme is 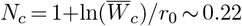.

Thus, when demes are at a small fraction (here, ~22%) of carrying capacity, 0.75 migrants per deme would still require a rather high rate of migration per individual– in this example, *Km*_*c*_ =0.75*/*0.22~ 3.4 (fig. 4c), which translates into 3.4 migrants per generation between demes at carrying capacity.

Figure 4c shows that the critical migration threshold *Km*_*c*_ always increases with increasing (2*Lu*)*/r*_0_, regardless of *h* and *Ks*. Moreover, the critical number *N*_*c*_*Km*_*c*_ of migrants per deme that is required to avoid extinction also increases with (2*Lu*)*/r*_0_, despite the fact that the (scaled) population size per deme *N*_*c*_ at the extinction threshold is lower at higher (2*Lu*)*/r*_0_. In other words, as selection becomes increasingly hard (either due to a higher genomewide deleterious mutation rate or lower baseline growth rate or both), each deme must exchange a larger number of migrants with other demes to avoid extinction, despite there being fewer individuals (that can potentially migrate) per deme.

Further, fig. 4e shows that − ln 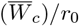 (the scaled load in populations at the threshold of extinction) also increases with (2*Lu*)*/r*_0_. However, this is not a simple linear increase, as one would expect if load were unaffected by genetic drift and determined solely by the total mutation rate. Instead, we find that for low values of (2*Lu*)*/r*_0_, load is significantly higher than the deterministic expectation 2*Lu*, suggesting that populations can remain stable despite a moderate inflation of load due to drift, as long as selection is not very hard. By contrast, with very hard selection, i.e., (2*Lu*)*/r*_0_ close to 1, load needs to be at its minimum possible value for populations to avoid extinction. For additive or weakly recessive alleles (1*/*2*<h<*1*/*4), this minimum is 2*Lu* (the deterministic expectation), while for highly recessive alleles, load can be significantly less than 2*Lu* due to purging.

### Load in non-extinct populations

Having analysed extinction thresholds, we now consider the *Km>Km*_*c*_ regime, where populations are stable. In this parameter regime, population sizes depend only weakly on *Km*, especially when alleles are recessive (fig. 3a), with load close to the deterministic prediction above *Km*_*c*_ (fig. 5b). This is contrary to previous work (Whitlock, 2002), which suggests that lower gene flow would reduce load under hard selection by allowing for more effective purging of deleterious alleles. To better understand our results, we plot the equilibrium allele frequency 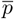 and mean load –ln 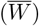 in the metapopulation as a function of *Km*, scaling these respectively by *p*_*det*_ and –ln 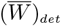, the corresponding (deterministic) allele frequency and load in a large undivided population (figs. 5a and 5b). The solid lines are results from the semi-deterministic approximation and colored circles represent simulation results. Here, we have obtained the deterministic *p*_*det*_ by solving −*sp*_*det*_(1−*p*_*det*_)(*h*+(1−2*h*)*p*_*det*_)+*u*(1−2*p*_*det*_)=0 and then using this to compute ln 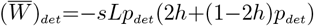; this is as opposed to assuming *p*_*det*_~*u/hs* and –ln 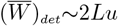, which are overestimates when alleles are nearly fully recessive.

We see from fig. 5a that when alleles are (partially) recessive, there is a decrease in the frequency of the deleterious allele (as migration decreases) due to purging. However, fig. 5b shows that this purging effect is not strong enough to counter the increased inbreeding at lower migration rates. Thus, while increased purging does cause load to be slightly lower than the deterministic expectation at intermediate migration levels and for highly recessive effects (*h*=0.02), this is not a significant effect. Similar figures for strongly selected alleles (*Ks*=10) are shown in Online Supplement E.

Thus, on the whole, reduced migration is detrimental to populations, with load increasing as populations become more isolated, until it exceeds a critical threshold ln 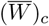 (which depends on the total mutation target and effect sizes of deleterious alleles; see fig. 4e), beyond which populations collapse. Importantly, neglecting the fact that both purging and fixation of deleterious alleles depend on local deme sizes (which shrink with falling migration due to eco-evo feedbacks) can mislead, causing us to overestimate the extent to which load is purged by greater expression of homozygotes in populations with limited migration.

## Discussion

This study explores the varied effects of gene flow, selection and population growth on load and extinction in a metapopulation. We distinguish between two modes of selection: soft selection, which applies when total load is much less than the baseline growth rate (i.e., (2*Lu*)*/r*_0_≪1), so that subpopulations are always at carrying capacity; and hard selection (with (2*Lu*)*/r*_0_≲1), where population sizes decline with increasing load (typically due to changes in allele frequency across multiple loci), which in turn, further increases load (due to less efficient selection in smaller populations). Our results thus clarify how eco-evolutionary feedback shapes the genetic diversity and long-term persistence of fragmented populations.

It has long been recognized that the reduction in mean population fitness due to drift load, i.e., the stochastic fixation of weakly deleterious (*Ks*~1) mutations, greatly exceeds the reduction due to segregating strongly deleterious mutations that are essentially maintained at deterministic mutation-selection balance (Kimura et al., 1963; Kondrashov, 1995). Our results suggests that as long as selection is soft, populations have to be very strongly fragmented in order for drift load to be a real issue. As shown in fig. 1a, very little migration (as little as one migrant per ~10 generations, *Km*=0.1) between a set of interconnected patches is enough to reduce the load due to slightly deleterious mutations (i.e., those with *Ks<*1) by a factor of 100, bringing it close to the deterministic expectation, regardless of dominance coefficient.

At higher migration rates, load is purely due to segregating (rather than fixed) alleles, with selection against recessive alleles being most efficient at intermediate *Km*. This is consistent with previous work that shows that load due to partially recessive alleles is lowest when migration is strong enough to prevent local fixation of deleterious alleles but not so high as to prevent efficient purging of recessives (Roze (2015); Whitlock (2002); Zhou and Pannell (2010)). Here, we further expand previous theory on soft selection by developing two analytical approximations that describe the evolution of moderately deleterious alleles under low to moderate levels of gene flow. These are: the ‘moderate selection’ approximation (lines in figs. 1a, 1c and 1d) that accurately captures how modest levels of selection (0.5*<Ks<*5) reduce local fixations (and thus *F*_*ST*_) at individual loci, and an approximation based on effective migration rates that captures the effects of multi-locus heterosis on allele frequencies and load (fig. 2).

Models of soft selection are relevant for two reasons. First, some fitness components affect population size more than others (adult viability versus male mating success, say). Second, estimates of load under the soft selection model can be an upper bound to the load in real populations, answering the following question: if the effective population size stays the same, what fraction of offspring will fail to survive under the predicted burden of deleterious mutations?

However, in reality, the accumulation of deleterious mutations will often decrease the size of the population, even leading to extinction (Kondrashov, 1995). Soft selection models do not take into account the positive feedback that ensues when the population size decreases due to genetic load, which in turn further increases genetic load (via increased drift), necessitating models that explicitly account for the feedback between evolution and demography. Such feedback has been investigated in earlier work on mutational meltdown (e.g., Lynch et al. 1995a,b), albeit only using simulations. Here, we develop a more comprehensive understanding using a semi-deterministic analysis which accounts for the stochastic effects of genetic drift on allele frequencies but neglects those of demographic stochasticity (Sachdeva et al., 2022).

A key result is that under hard selection, demes must exchange a much higher number of migrants per generation despite supporting fewer individuals (figs. 4c and 4d)– this requires very high rates of migration per individual. Further, critical migration thresholds *Km*_*c*_ are highly sensitive to (2*Lu*)*/r*_0_ (a proxy for the hardness of selection), with higher values of (2*Lu*)*/r*_0_ necessitating more gene flow. This is because when the total deleterious mutation rate (and consequently, mutation load) is comparable to the baseline growth rate, then the eco-evo feedback between population size and load is strong enough to amplify any fluctuations in load (e.g., due to slightly reduced gene flow and/or increased drift). Very high levels of migration are then required to buffer individual demes against abrupt collapse and prevent global extinction.

The parameter (2*Lu*)*/r*_0_, which emerges as a crucial parameter in our analysis, captures the extent to which deleterious mutations (occurring at a genomewide rate 2*Lu*) reduce the baseline growth rate *r*_0_. Estimating (2*Lu*)*/r*_0_ thus requires knowledge of both the total deleterious mutation rate (e.g., from mutation accumulation experiments or genomic methods) and the baseline growth rate *r*_0_ of a maximally fit reference population. However, *r*_0_ is difficult to measure or even define unambiguously— for instance, one could define it either via the reproductive output of the ‘least loaded’ class in a specific population, or as the rate of growth associated with a hypothetical, perfectly fit genotype, with the former being (possibly) easier to measure but harder to generalise across populations. Another source of complexity is that deleterious mutation rates (or more generally, the amount of resources allocated to germline maintenance) may co-evolve with growth rates due to trade-offs between the two (Avila and Lehmann, 2023), so that composite parameters such as (2*Lu*)*/r*_0_ are somewhat constrained.

We also explore how the critical migration threshold *Km*_*c*_ for metapopulation persistence depends on selection and dominance coefficients of individual alleles, and find that *Km*_*c*_ (as well as *N*_*c*_*Km*_*c*_, the number of migrants exchanged per deme) is highest for intermediate values of *Ks* (~1) and for additive (as opposed to recessive) alleles (fig. 4). These results mirror those under soft selection, where load is highest for intermediate *Ks* and *h*~1*/*2 (fig. 1). The finding that populations can survive a higher degree of fragmentation (i.e., lower levels of gene flow) when load is primarily recessive appears to be at odds with verbal arguments that draw a (somewhat loose) connection between fragmentation and increased inbreeding load due to expression of recessives. Our analysis refines this argument by showing that when load is largely due to recessive alleles, mutational meltdown is triggered more ‘easily’ in the sense that population sizes per deme are relatively high (and load low) at the extinction threshold (fig. 4e). However, mutational meltdown is also more ‘difficult’ in the sense that populations can survive with lower levels of migration (without accumulating too much load), than if load were primarily additive.

Our stochastic (finite *K*) simulations show that close to the critical extinction threshold, local deme sizes follow a bimodal distribution (figs. 3c and 3d), with some demes teetering on the brink of extinction while others support stable populations. Such bimodal distributions (and more generally, any differences across demes) arise due to the stochastic nature of the model, despite the fact that growth rates, carrying capacities and selection coefficients are uniform across demes. Such bimodality can serve as an important signal of impending population collapse, where minor variations in migration rate or external factors can push some demes past the point of no return, causing local extinctions and a possible cascade towards global extinction (Drake and Griffen, 2010; Scheffer et al., 2001).

Our findings, while intuitive, can become obscured by the confusion around ‘hard selection’. In the context of subdivided populations, this term is used to describe scenarios where the genetic contribution of any sub-population to the next generation is proportional to its fitness. However, this can encompass two fundamentally different kinds of models. In the first kind, the size of the full population (as well as of sub-populations) is determined by global density-dependent regulation, and thus does not change from one generation to the next (Christiansen, 1975; Whitlock, 2002)– an assumption that precludes mutational meltdown and extinction. By contrast, in the second kind of model, the size of each sub-population is determined by its local (fitness-dependent) growth rate and local density-dependent regulation (Higgins and Lynch, 2001; Polechová and Barton, 2015; Szép et al., 2021). In these models, not only do less fit populations send out fewer migrants, but can also shrink in size, accumulating more deleterious mutations as they do, which further aggravates population decline, potentially leading to extinction. These two kinds of models– both ostensibly allowing for ‘hard selection’ (in that low-fitness populations contribute less to future generations)– yield qualitatively different predictions. For example, we have shown that reduced gene flow is largely detrimental to populations– increasing drift load and the risk of extinction. This is in contrast to previous work on hard selection in subdivided populations (Whitlock, 2002), which finds that reduced gene flow always *decreases* load (via greater expression of homozygotes and more efficient purging), but only while neglecting the dependence of (local or global) population sizes on fitness.

Our analytical approach is based on the continuous time limit, which applies when ecological and evolutionary processes are sufficiently slow. In particular, this requires population growth rates to be small, which may not hold in general (e.g., during rapid population expansion), but is reasonable for populations close to equilibrium. In the continuous time limit, the distribution of allele frequencies and population sizes is *independent* of lifecycle details and can be described via the diffusion approximation, which depends only on scaled parameters (see eqs. (2) and (3)). The diffusion approximation is a cornerstone of population genetics (Kimura, 1955), though much less used in ecology and the study of eco-evolutionary feedback. Our study demonstrates that at least in some regimes, theory formulated in terms of continuous time is a powerful alternative to analysing eco-evo feedback in terms of selection gradients that account for lifecycle details– an approach that is more common in the field (Gandon and Michalakis, 1999; Parvinen et al., 2020; Rousset and Ronce, 2004). However, in contrast to our study, these analyses usually assume that relatedness at genetic loci influencing ecological traits is unaffected by selection– a very strong assumption that only holds if *Ks*≪1, i.e., if local drift is much stronger than selection per locus (inset, fig. 1d).

What are some limitations of our study? Our theoretical framework is based on the infinite-island model, which neglects explicit space. Although this is a useful simplification that ensures analytical tractability, in reality, spatial structure can strongly affect metapopulation dynamics. For instance, Bascompte and Solé (1996) argued that metapopulations embedded in space are more vulnerable to extinction than would be suggested by spatially implicit (e.g., island) models – thus, the latter may underestimate critical thresholds and the impact of fragmentation on metapopulation persistence. This begs the question: are there specific spatial configurations more conducive to the maintenance of genetic diversity, and how can spatially and genetically explicit models inform practical decisions, e.g., the planning of migration corridors (Bennett, 1999) or assisted gene flow (Aitken and Whitlock, 2013) across continuous space?

Our study also does not account for the dynamic, context-dependent nature of migration (Ronce, 2007). For instance, dispersal-related traits are often quite heritable (Saastamoinen et al., 2018), making it possible that dispersal rates might evolve (or change in a plastic manner) in response to increased fragmentation and kin competition. The evolution of dispersal has been modeled in several studies, which investigate evolutionary stable levels of migration under a range of demographic scenarios (Gandon and Michalakis, 1999; McPeek and Holt, 1992; Parvinen et al., 2020), often focusing on the inclusive fitness of dispersal-altering alleles (Rousset and Ronce, 2004). However, this work largely ignores the genetic architecture of inbreeding load or heterosis (though see Roze and Rousset, 2005), which may be important drivers of dispersal evolution. More generally, given observations of phenotypic (and possibly, genetic) correlation between traits that affect dispersal and those affecting fitness (Saastamoinen et al., 2018), a key question is how dispersal, load and population sizes might co-evolve, e.g., in the presence of trade-offs between fitness and dispersal, with such trade-offs potentially generating balancing selection on alleles that affect both. Extending theoretical frameworks that explicitly model the genetics of load (such as the one here or in Roze and Rousset, 2005) to these more realistic scenarios is an interesting direction for future work.

Our study, while accounting for the polygenic nature of load, does nevertheless make various simplifying assumptions about its genetics. First, we neglect epistatic interactions between loci. Negative epistasis, in particular, increases the efficiency of selection against deleterious alleles (Kimura and Maruyama, 1966), though this effect has been argued to be too weak to resolve the paradox of drift load (Charlesworth, 2013). Second, we assume that deleterious alleles are unlinked, so that Hill-Robertson interference within populations can be neglected. How might this change with lower rates of recombination or in specific genomic regions with arrested recombination, e.g., inversions? On the one hand, reduced recombination should exacerbate drift load by making selection less efficient; on the other hand, multi-locus heterosis may be stronger (Berdan et al., 2021). Third, we have considered spatially uniform selection. It would be interesting to extend our framework to scenarios where some loci are under spatially heterogeneous selection while others are subject to unconditionally deleterious mutation. Gene flow may then swamp local adaptation, while concurrently alleviating the burden of globally deleterious mutations. Determining the relative strengths of these contrasting effects under realistic parameter regimes is crucial for understanding adaptation and extinction in fragmented, heterogeneous landscapes.

## Acknowledgements

This research was partially funded by the Austrian Science Fund (FWF) [FWF P-32896B] and DOC Fellowships of the Austrian Academy of Sciences: grants 26380 (O. Olusanya) and 26293 (K. Khudiakova). We thank Nick Barton for useful comments on the thesis chapter that led to this manuscript.

## Statement of Authorship

Conceptualization and Model analysis: O.O, K.K and H.S; Coding simulation: O.O and H.S; Supervision: H.S; Writing (original draft): O.O, K.S and H.S; Writing (review and editing): O.O and H.S.

## Data and Code Accessibility

The Fortran and Mathematica codes and simulations to reproduce our analyses are available online at https://zenodo.org/records/14682599 (DOI: https://doi.org/10.5281/zenodo.14682599).

## Supplement A. Details of allele frequency simulations

We assume a life cycle consisting of migration between patches, mutation, followed by viability selection on adults.

To account for the effect of migration on population size, we first determine the net number of migrants, *netM* (*t*) (i.e., the difference between the total number of immigrants and emigrants in a given patch) and add this to the existing patch size. Under soft selection, we assume a zero net migration rate (i.e., a balance between the number of immigrants and emigrants) so as to keep the patch size fixed. However, under hard selection, the net number of migrant is assumed to be non-negative. Migration therefore changes the population size according to 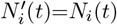 (under soft selection) and 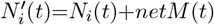 (under hard selection). Similarly, to account for the effect of migration on allele frequencies in any patch, we add the net number of migrant alleles, *netp*(*t*) to the existing allele copy number in the patch and divide this by the population size 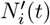. In essence, migration changes allele frequencies according to 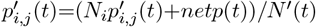.

With regards to the effect of mutation, we assume equal mutation rate to and from the deleterious allele so that mutation changes allele frequency according to 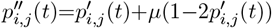. However, mutation has no effect on population size (under hard selection) so that 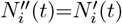.

Following mutation, adults in each patch mate to produce offspring that survive to be next generation parents. Under soft selection, the mating process in each patch involves sampling with replacement, *N*_*i*_(*t*) pair of individuals based on their fitness and freely recombining their gametes to form offspring gametes. This means that we do not explicitly distinguish between the male and female sexes and there is also the possibility of self mating. Under hard selection on the other hand, load and density-dependent regulation changes population size according to 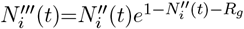 and demographic stochasticity (randomness in birth and death) is further imposed on the population to determine how much survivable offspring, 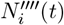 can be formed. The latter is achieved by sampling from a Poisson distribution with parameter 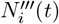. Offspring are then formed by choosing with replacement 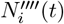 pairs of parents also based on their fitness and freely recombining their gametes. For both soft and hard selection, selection changes allele frequency according to 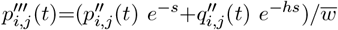 where 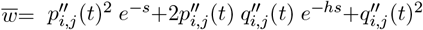 is the mean fitness in the *i*^*th*^ patch and 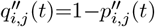.

Finally, the allele frequency at the end of the generation, i.e., 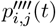 is obtained by sampling 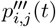 from a Binomial distribution with parameters, 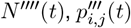 thus accounting for genetic drift.

## Supplement B. ‘Moderate selection’ approximation under soft selection

Under the soft selection model, any quantity such as load, *F*_*ST*_, etc. for a single locus can be computed from the equilibrium allele frequency distribution 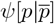, by first numerically solving 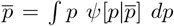 to obtain the mean allele frequency 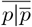, then plugging this into the equilibrium distribution 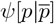, and finally integrating over *ψ*[*p*] to obtain higher moments. However, it is also useful to consider an alternative approach based on moments (or cumulants) of *ψ*[*p*].

In general, any moment (such as 𝔼[*p*] or 𝔼[*p*^2^]) will depend on higher moments, resulting in a set of recursions that is not closed. Approximations thus rely on closing this set of recursions in different ways, depending on assumptions about the relative magnitudes of *Ks* (or *Ksh*) and *Km* (Whitlock (2002); Glémin et al. (2003); Roze (2015)). Here, we introduce a new moment closure approximation, which applies also for recessive (*h*~0) alleles and intermediate selection coefficients (*Ks*~1).

As in the main text, let *p* denote the frequency of the deleterious allele at a given locus in a given deme and 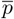 the mean across all demes in the population. We denote the expected change in allele frequency per unit time by *M* (*p*) and the variance of the change by *V* (*p*). These are:

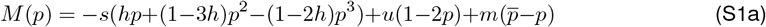

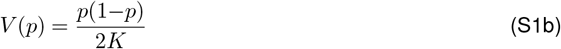

where *K* is the number of individuals per deme. Let 𝔼_*ψ*_[*f* (*p*)] denote the expectation of any function *f* (*p*) of the allele frequency *p* over the frequency distribution *ψ*[*p*]. Under the diffusion approximation, 𝔼_*ψ*_[*f* (*p*)] satisfies (see also Ohta and Kimura (1971)):

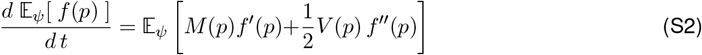

Setting *f* (*p*) as *p* and *p*^2^ yields the following equations for the first and second moments of the allele frequency distribution respectively (see also Whitlock (2002), Glémin et al. (2003)):

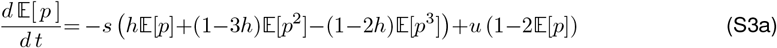

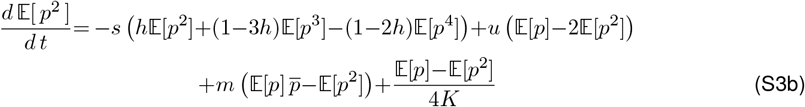

It will be useful to express the above equations in terms of appropriately scaled cumulants of the frequency distribution:

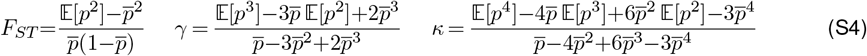

Here, *F*_*ST*_, *γ* and *κ* denote respectively the variance, skew and kurtosis of the frequency distribution within any deme, scaled by the corresponding cumulant for the distribution of frequencies across all demes.

At equilibrium, moments of the frequency distribution are constant in time, i.e., *d* 𝔼[*p*]*/dt* = *d* 𝔼[*p*^2^]*/dt* = 0; further, 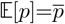. Combining equations (S3) and (S4), and assuming deleterious alleles to be sufficiently rare *overall* in the population that 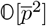 terms can be neglected, we have at equilibrium:

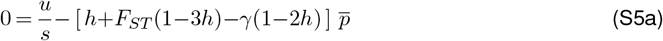

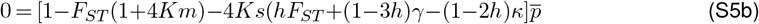

This *pair* of equations is under-determined as it involves *four* variables 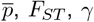, and *κ*. To obtain an approximate solution, we assume further that third and higher cumulants i.e., the skew and kurtosis are related to *F*_*ST*_ as in the *neutral* infinite-island model. Note that *F*_*ST*_ itself is not taken to be neutral (or unaffected by selection), as is assumed by Whitlock (2002), but only that:

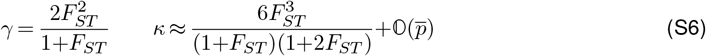

Substituting into eq. (S5) and expressing load as 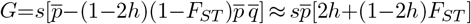, we obtain:

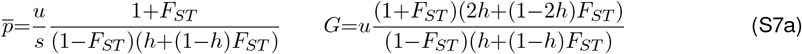

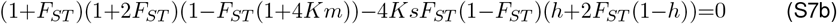

Equation (S7b) is cubic in *F*_*ST*_ and can be solved (e.g., numerically); this solution for *F*_*ST*_ can then be sub-stituted into eq. (S7a) to obtain the average deleterious allele frequency 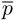 and load *G*. Thus, in essence, our approximation allows for selection that is strong enough to change *F*_*ST*_ but not the relationship between higher cumulants (*γ, κ* etc.) and *F*_*ST*_.

It is useful to juxtapose this approximation with those used in earlier work. Whitlock (2002) assumes *weak selection* such that *F*_*ST*_ is essentially neutral, i.e., equal to 1*/*(1+4*Km*). This is valid when *Ks*≪ 1, so that the term involving *Ks* in eq. (S7b) (or equivalently in eq. (S5b)) can be neglected. At the other extreme, Roze (2015) and Glémin et al. (2003) consider a parameter regime where the local deme size *K* is large enough that allele frequency distributions are essentially concentrated around the mean 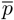, which is relatively low. In practice, this means that *F*_*ST*_ is small enough that 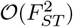 terms in eq. (S7b) can be neglected, which gives: 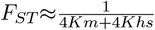. Theirs is thus a *strong selection, strong migration* approximation; in practice, it applies for *Ksh*+*Km* greater than 5. By contrast, the approximation for *F*_*ST*_ introduced above applies also for intermediate *Ks* (see fig. 1d in the main text) and is thus a *moderate selection* approximation: in essence, it captures the key effect of selection which is to change coalescence times within and between demes (to *different* extents) without necessarily changing the extent to which the branching structure of genealogies is (a-)symmetric. However, this is only a rough interpretation, and more rigorous analyses are required to understand intermediate *Ks* regimes where neither selection nor drift can be treated as minor perturbations (to neutral or deterministic predictions respectively).

### Relaxing the assumption of 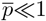

All the approximations described above assume that 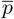 is sufficiently low (i..e, deleterious alleles so far from global fixation) that 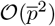 terms can be neglected. When this is no longer true (but if (S6) still applies, i.e., if third and higher cumulants depend on *F*_*ST*_ as in the neutral model), equations (S3), (S4) and (S6) together yield a cubic equation for 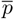:

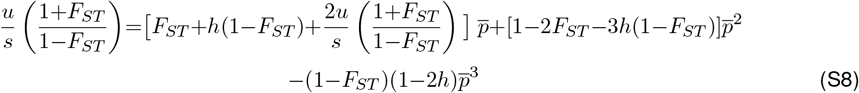

Since deleterious allele frequencies are expected to be high only for very weak selection, we can typically neglect the effects of selection on *F*_*ST*_ in this parameter regime, and simply solve equation (S8) for 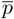 under the assumption 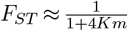.

## Supplement C. Extinction thresholds with a distribution of selection and dominance coefficients

Here, we explore the impact of gene flow on the mean population size in the metapopulation when we have a distribution of selection and dominance coefficients. We only consider a simple distribution, where a fraction *a* of loci are additive (*h*=0.5) and have scaled selection coefficients *Ks*_*A*_ and the remaining fraction, (1−*a*) are nearly recessive (*h*=0.02), with selection coefficients *Ks*_*R*_.

We see from fig. S1a - S1c that independent of the value of *a*, when additive alleles are nearly neutral and recessive alleles are non-neutral, increasing the selective effect of the recessive alleles, i.e., making them more strongly deleterious has little or no effect on the critical migration threshold above which the metapopulation survives. This also holds for the case where the additive alleles are mildly deleterious and occupy a higher proportion of loci (dashed lines in fig. S1c).

**Figure S1:**
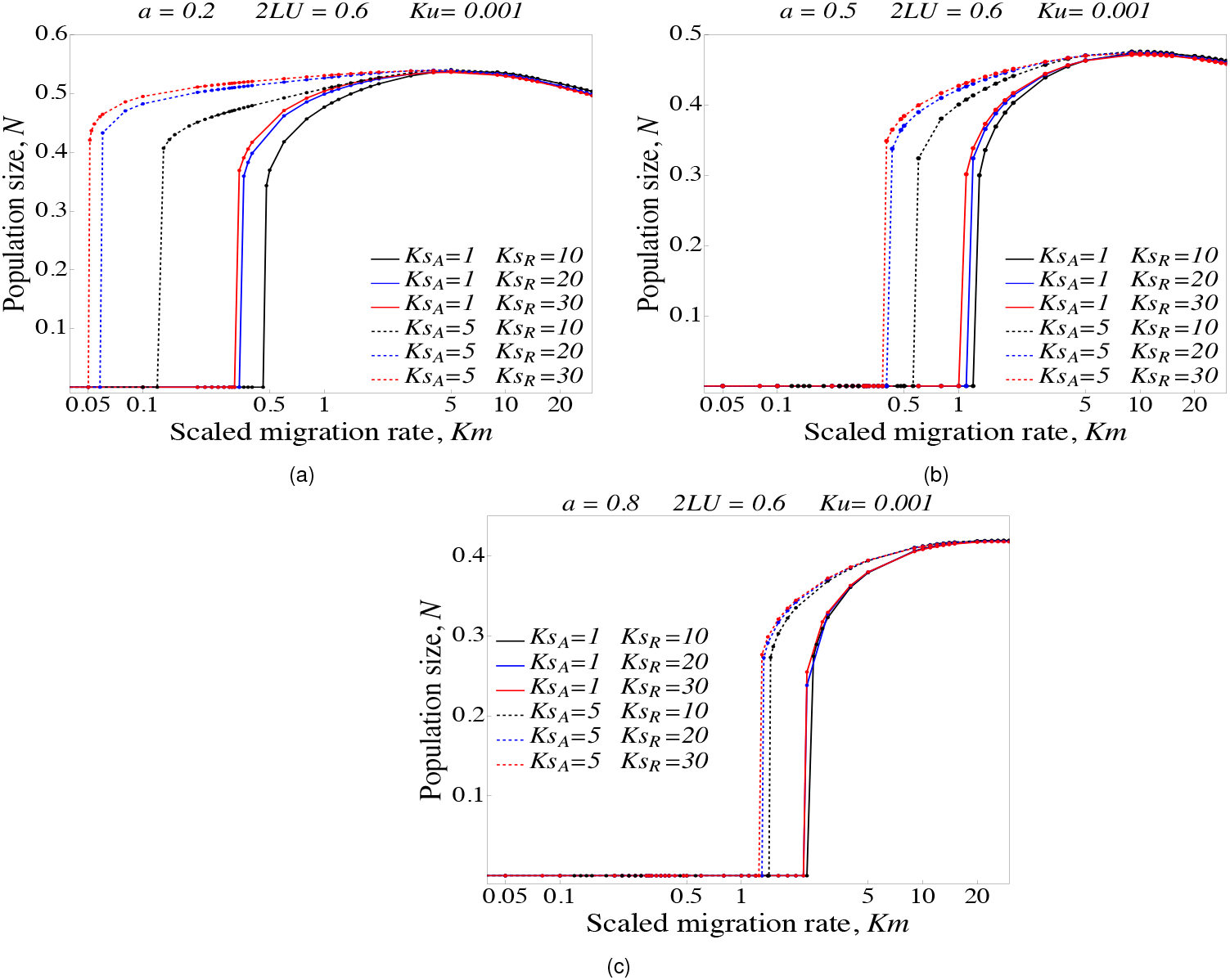
Average population size across the metapopulation plotted against *Km* with (a.) *a*=0.2 i.e., 20% of loci are additive and 80% are recessive (b.) *a*=0.5 i.e., with equal proportion of additive and recessive alleles (c.) *a*=0.8 i.e., with 80% of loci being additive and 20% recessive.

On the other hand, when additive alleles occupy a lower or equal proportion of loci as the recessive alleles, we see a somewhat different dynamics. We observe a (slightly) higher critical migration threshold when additive and recessive alleles are mildly deleterious (black dashed line in fig. S1a and S1b). As the recessive alleles become moderately deleterious, the threshold reduces and increasing the selective effect further makes little or no effect on the threshold migration rate (compare blue and red dashed lines in both fig. S1b and S1c). Finally, for all *Ks*_*A*_, *Ks*_*R*_ combinations considered, we see the critical migration threshold at least doubling as *a* increases.

## Supplement D. Comparison of semi-deterministic predictions with simulations (*h*=1*/*2)

**Figure S2:**
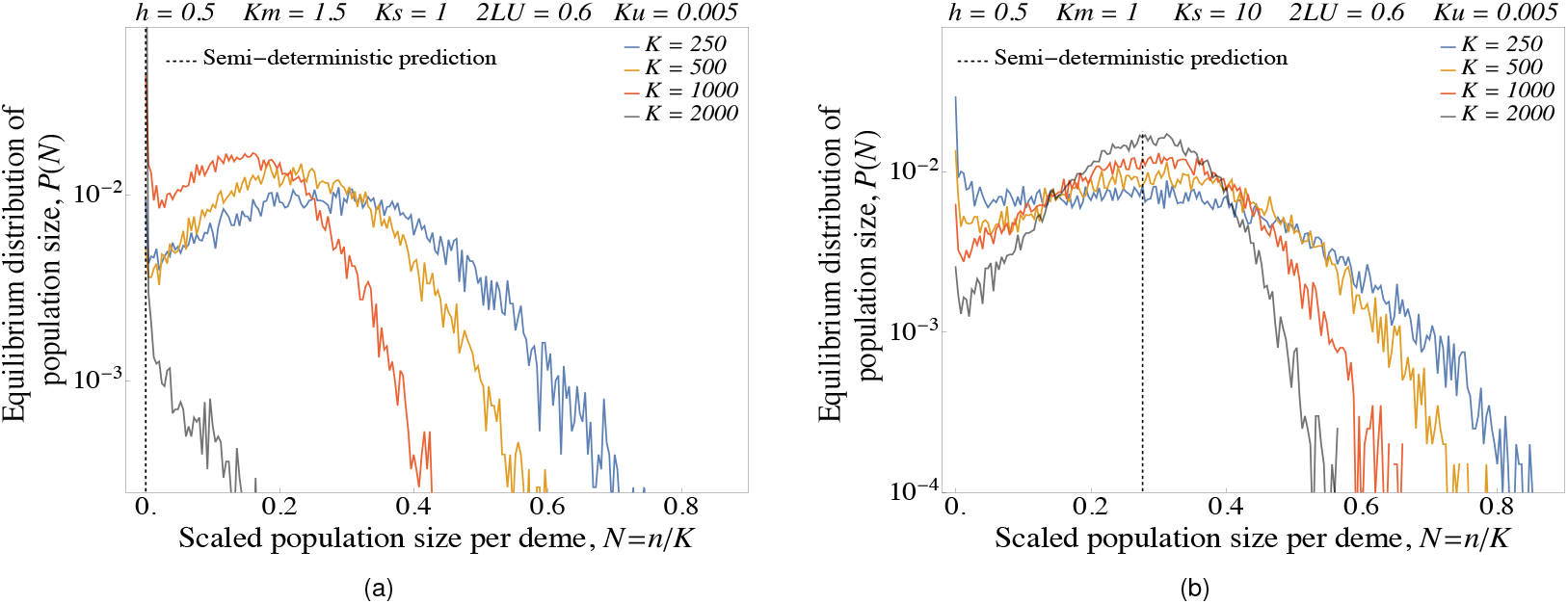
Comparison of simulation results with semi-deterministic predictions when load is due to additive alleles. Figures show equilibrium distribution *P* (*N*) of population sizes *N*, as obtained from simulations, for various values of *K*, for two different scaled parameter combinations for which the semi-deterministic approximation predicts extinction (in (a)) vs. stable populations (in (b)). The scaled parameters *Ks, Ku, Km*, 2*LU* =2*L*(*u/r*_0_) are held constant as *K* is varied (by simultaneously varying *s, u, L*, and *m*). The scaled parameters have values: (c) *Ks*=1, *Km*=1.5, or (d) *Ks*=10, *Km*=1, with other parameters being *h*=0.5, *Ku*=0.005 and 2*LU* =0.6, as above. Black vertical dashed lines represent the semi-deterministic prediction for *N* : this is *N* =0 in (c) and *N* =0.276 in (d). The distribution *P* (*N*) becomes sharply peaked around the semi-deterministic prediction as *K* increases in both (c) and (d). All simulations were performed with 100 demes and *r*_0_=0.1.

Here, we demonstrate the convergence of simulation results to the semi-deterministic prediction with increasing *K* more generally, by considering load due to additive (*h*=1*/*2) alleles, in contrast to figures 3c and 3d in the main text, which show results for nearly recessive (*h*=0.02) alleles. As in the main text, we consider two different parameter combinations for which the semi-deterministic approximation predicts global extinction (fig. S2a) vs. stable populations with *N*_*_=0.276 (fig. S2b). As in the case of recessive alleles (figures 3c and 3d in the main text), we find that the distribution of scaled population sizes *P* (*N*) becomes narrower with increasing *K* (which corresponds to weakening demographic fluctuations). In fig. S2a, this causes the mean population size to decrease (since there are fewer large islands that can act as demographic sources), with the entire population nearly extinct at *K*=2000 (as also predicted by the semi-deterministic analysis). By contrast, in fig. S2b the distribution of deme sizes gets concentrated around a finite *N*_*_ (which matches the semi-deterministic prediction) with increasing *K*.

## Supplement E. Effect of gene flow on equilibrium allele frequency and equilibrium mean load for *Ks*=10

**Figure S3:**
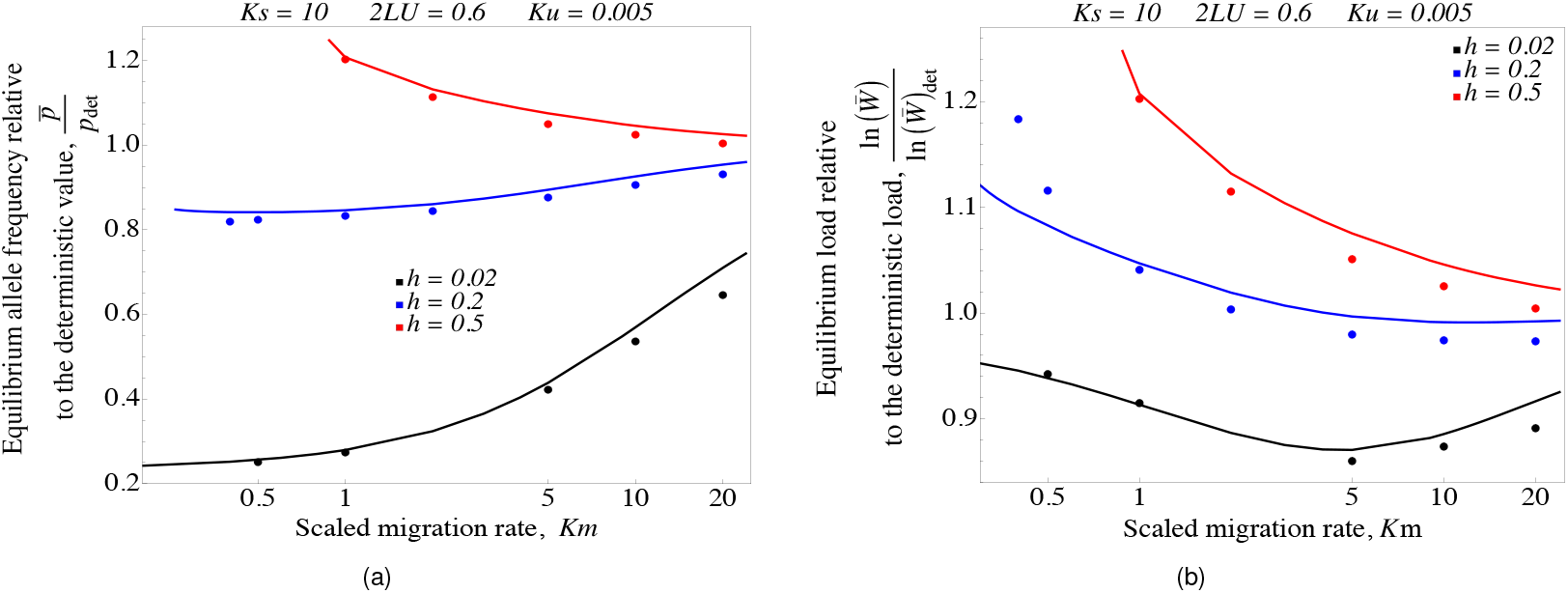
(a) Equilibrium frequency 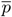 of deleterious alleles and (b) equilibrium load *R*_*g*_ (relative to *p*_*det*_ and *R*_*g,det*_, the deterministic allele frequency and load in a large undivided population), plotted against *Km*, for *Km>Km*_*c*_. Filled circles show simulation results (using allele frequency simulations) and solid lines show results from the semi-deterministic approximation. The scaled parameters are *Ku*=0.001, *Ks*=1 and 2*LU* =0.6; simulations are run with 100 demes with *K*=3000 and *r*_0_=0.1.

## Notes

https://doi.org/10.5281/zenodo.14682599

### Competing Interest Statement

The authors have declared no competing interest.

### Summary of Updates

This is an updated version of the manuscript.

